# Creation of genomic chimeras via megabase scale genome expansion

**DOI:** 10.64898/2026.11.04.764350

**Authors:** Charles Sanfiorenzo, Raymond J. Zhang, George Wythes, Megan Wang, Jianyi Huang, Jeongsoo Hur, Mikel Lipschitz, Jie Zhou, Russell Swift, Bryan Gerber, Kaihang Wang

## Abstract

An organism’s physiology is controlled by its genomic content. Building genomic chimeras would enable melding of the diverse functions and properties of life. However, prior arts in genome synthesis are limited to reconstituting pre-existing functions within a singular genome rather than combining diverse genomic functions across multiple distinct genomes. Existing methods are also prohibitively expensive, labor intensive, and time consuming, and thus fundamentally not scalable for creating large synthetic chimeric genomes. To address all these limitations, we invented Additive Conjugative-CAST Engineering (ACE) combining conjugation with CRISPR-associated transposition (CAST) to deliver and integrate up to half a genome per step from a donor cell into a precisely defined position in the recipient’s genome. We have demonstrated ACE’s capability to integrate at least a 2-megabase donor genomic segment in a single step and at least 3 megabase in two consecutive steps. Importantly, we confirmed ACE’s generality through the creation of multiple genomic chimeras across species, genus, and at least order barriers. With ACE, we further demonstrate such chimeric organisms, or what we call genome expanded organisms (GEOs), can be stably forged from at least three starting bacterial strains, and can be stably maintained to express all their acquired genomic parts in a single chimera. ACE offers a new paradigm of creating artificial lifeforms to combine desired properties and potentially create novel functions beyond the constraints of nature. ACE also offers a unique perspective to probe unexplored plasticity in genome architecture and expression patterns of novel, massively expanded genomes.

## Main

The design and construction of artificial genomes furthers our ability to understand and engineer biology^1,2,3,4,5,6^,. Organisms with designer genomes allow us to disentangle the complexities of gene functionality and essentiality^7,8^, transcriptional and translational control^9,10^, and genome fold and architecture^11,12^. Moreover, artificial genomes enable us to engineer novel properties and encode new functions into synthetic organisms^13,14^, enhancing the potential of living systems to produce artificial products and perform programmed tasks beyond the constraints of nature^15,16^.

Important benchmarks in this emerging field include the assembly of minimized *Mycoplasma* genomes^7,17,18^, the recoding of *Escherichia coli* Syn61 genome^9,14^, and the recent consolidation of multiple synthetic chromosomes in the “*S. cerevisiae* 2.0” synthetic yeast genome project,^8,19^. On-going efforts include the *E. coli* Syn57 project^20^, and the *Salmonella typhimurium* recoding project^21^. In general, their workflows all follow a similar paradigm consisting of four steps: (1) *in silico* design of the synthetic genome, (2) synthesis and assembly of synthetic DNA (syn. DNA), (3) delivery of assembled syn. DNA into target cells, (4) replacement of corresponding genomic sequences with delivered syn. DNA^7,8,14^. Despite immense progress over the past two decades, fundamental challenges within these four steps make genome synthesis prohibitively expensive, labor intensive, and time consuming^2,6^.

Design of novel sequences for synthetic genomes is difficult. Because of the current inability in reliably predicting the function of *de novo* designed coding and non-coding DNA sequences for programmed artificial functions, all synthetic genome designs so far are constrained to reconstituting pre-existing natural functions by refactoring^8,14^, recoding^14,20^, minimization^7^, or rearrangement^8^ of existing sequences^3,5^. Despite this limited scope, seemingly small design flaws in such reconstituted genomic sequences can become major failure modes in a genome synthesis project^7,8,14,22^.

Genomic scale DNA assembly is also time, labor, and resource intensive. *De novo* oligo synthesis is currently limited to products 200-300 nucleotides in length by phosphoramidite chemistry and up to 1-kb by terminal deoxynucleotidyl transferase^2^. Through multiple rounds of hierarchical assembly, these short oligos are pieced together via 5 to 10-kb intermediates to 50 to 100-kb fragments for iterative genome replacements^8,9,14,19,23^ or even to 1-Mb complete genomes for genome transplantation^3^. Despite standardized automation in short oligo synthesis, DNA assembly is typically performed manually with limited throughput, and oftentimes require bespoke techniques for execution. Moreover, both the shorter intermediates and the assembled fragments need to be thoroughly sequenced in each step to eliminate the propagation of synthesis and amplification errors into the final construct. These factors have restricted synthetic genome endeavors to exceptionally well-funded institutions and consortia^6^.

Finally, the delivery of syn. DNA for the replacement of wildtype genomic sequence in living cells pose multiple challenges. Long DNA on the sub-megabase (> 100-kb) scale is prone to sheer stresses when handled *in vitro* and is difficult to deliver into living cells. Methods to introduce such large DNA into a living cell for subsequent genome integration are often specific for a particular chassis and have limited programmability. For instance, synthetic *Mycoplasma* construction involves genome transplantation techniques of near megabase sized constructs^24^. The *S. cerevisiae* 2.0 project revolves on intrinsic recombination to assemble and replace 50 to 100-kb syn. DNA into yeast chromosomes in each iterative step^23^. Both approaches have not been broadly generalizable to other organisms. Methods based on homologous recombination mediated by viral or bacterial host proteins such as genomic fission/fusion^25,26^, CAGE^27^, REXER^9^, and related methods^28^ have been pivotal in rearranging and building custom *E. coli* genomes but lack the ability to robustly streamline megabase-sized manipulation of highly homologous sequences and may not be applicable to other organisms.

Beyond replacement, there are currently even more limited solutions to perform large-scale genome insertions. The assembly of a fragmented *Synechocystis* genome across the *Bacillus subtilis* genome in multiple non-continuous segments using site-specific recombinase (SSRs) has suggested the possibility of building expanded designer genomes^29^. However, this work suffered from the challenges associated with the conventional genome construction workflow as the foreign genome was installed in 30-kb steps in *B. subtilis* over a decade and the inserted DNA was transcriptionally silent. As an alternative to stable genomic integration, addition of 0.5 to 1-Mb *E. coli* genome sequences into *E. coli* episomes with genomic fission or SSRs has been demonstrated^25,26,30^. Iterative additions using a REXER-based method has also been shown to assemble ∼1-Mb of inert human genomic sequence into an *E. coli* episome in nine consecutive steps^28^. However, such episomes require additional selection to maintain within the host bacterium and have a potentially limited carrying capacity beyond 1-Mb due to the intrinsic limitations of episomal replication and partitioning^26,30^. Both the requirement of stable, long-term maintenance and very large carrying capacity could be solved by integration into the host genome instead of the self-replicating episomes. Hence, a new genome construction technology that couples robust multi-megabase scale delivery with programmable multi-megabase scale genomic integration in a single step would introduce new versatility and power to the toolbox of artificial genome creation.

To address this need, we developed the Additive Conjugative-CAST Engineering (ACE) method. ACE combines an engineered conjugation module derived from the broad host range conjugative RP4 plasmid with an engineered Type I-F *Tn6677* CRISPR-associated transposition (CAST) module from *Vibrio cholerae* HE-45 to precisely transfer and insert defined donor genomic fragments into the recipient’s genome in a programmable manner^30,31,32^. CAST has been previously demonstrated to perform CRISPR RNA (crRNA) dependent DNA transposition of up to 10 kb into target genomic loci and function in a variety of bacterial species^33,34^. Inspired by its programmability and versatility, we adapt CAST as a part of ACE and expand its capacity to integrate massive DNA cargoes ranging from 100-kb to multi-megabase in a single step. We show efficient, precise, and robust ACE integrations of at least 2-Mb per step into the *E. coli* genome, and genome expansions of at least ∼3-Mb possible in two consecutive ACE steps. As a portable method, ACE functions across a wide host range spanning different genus and orders.

With ACE, we propose the concept of genome chimerism and the systematic creation of genome expanded organisms (GEOs) as a new paradigm to create artificial life with engineered functions and forms. We further demonstrate ACE as a tool for interrogating the genome architecture and the transcriptomic landscape of these chimeric cells with massive complementation of their DNA genome.

## Results

### Design, validation, and characterization of ACE

In the ACE system, the engineered bacterial conjugation and CAST components (**Figure 1**) are incorporated within four modular parts: Cassette^START^, Cassette^END^, *pAra-Trans*, and *pAra-CAST*. Cassette^START^ and Cassette^END^ each encode one origin of transfer (oriT) sequences from the RP4 conjugative plasmid and either the Right or Left transposon end (R-TE and L-TE respectively) sequences from the *Tn6677* CAST system, together with corresponding negative and positive selection markers and green fluorescent protein^35^ (**Figure 1a**). R-TE is placed downstream to the oriT in Cassette^START^ to mark the start of conjugation, while L-TE is placed upstream to the second oriT in Cassette^END^ to mark the end of conjugation. Together, Cassette^START^ and Cassette^END^ mark the precise start and the end of the transferred genomic fragment from the donor and retain the R-TE and L-TE sequences after the conjugation into the recipient cell to guide the genomic integration. Plasmid *pAra-Trans* holds the components of a minimized conjugation system to facilitate DNA conjugation from the donor cell (**Figure 1a**). Plasmid *pAra-CAST* encodes the rest of the CAST components, including the crRNA dictating the location of genomic integration in the recipient cell (**Figure 1a**).

**Figure 1.**
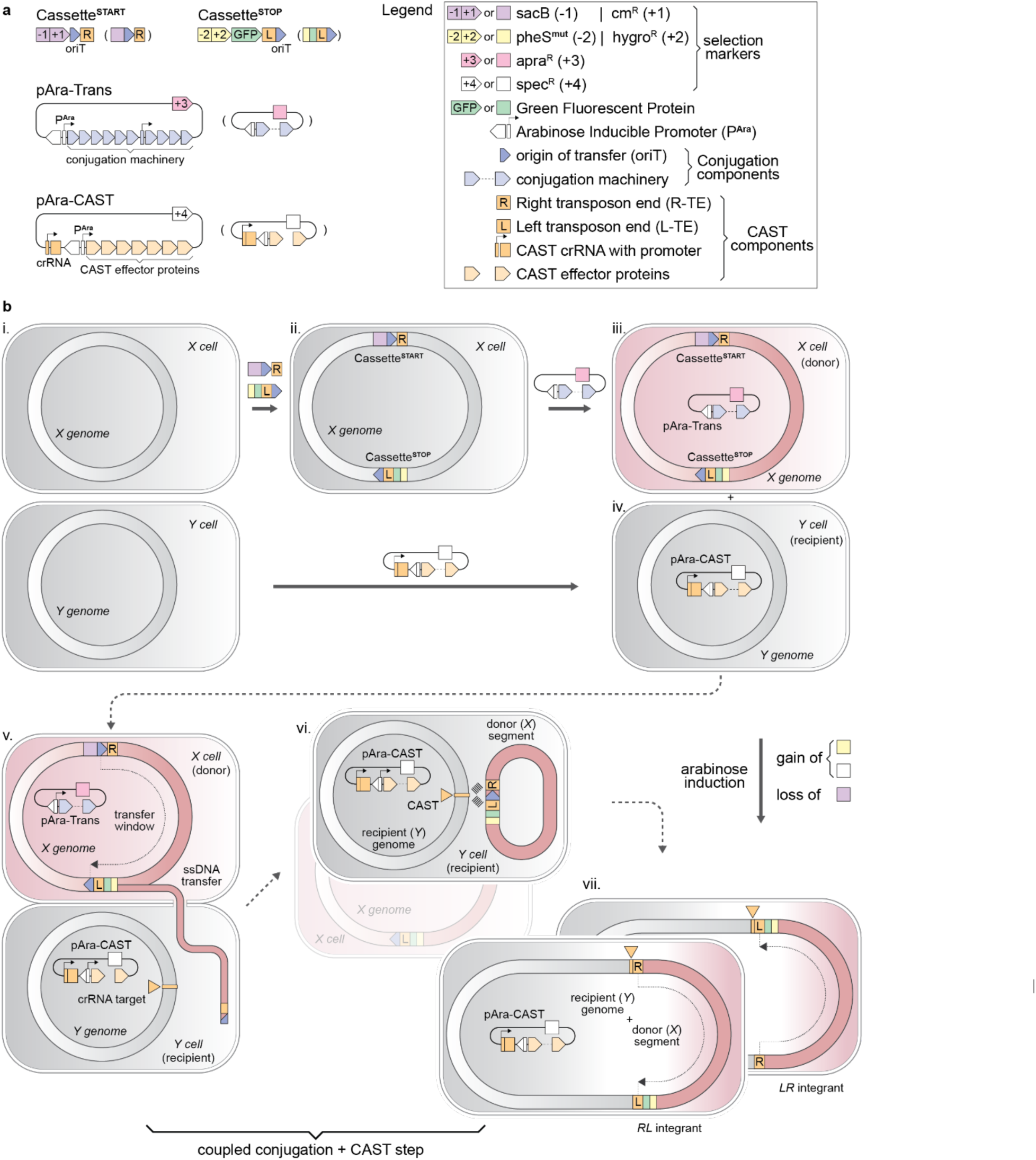
Design and overview of ACE. (**a**) Schematics of *Cassette^START^Cassette^END^* of ACE, together with designs of the *pAra-Trans* conjugation helper plasmid and the *pAra-CAST* transposition effector plasmid. *Cassette^START^*contains *sacB* (−1) and *cm^R^* (+1) genes encoding sucrose susceptibility and chloramphenicol resistance, respectively, in addition to a *RK24* origin of transfer (oriT) and the right end (RE) of the *Tn6677* transposable element. *Cassette^END^* contains *pheS* (−2) and *hygro^R^* (+2) genes encoding p-chloro-phenylalanine susceptibility and hygromycin resistance, respectively, in addition to a *gfp* encoding “Green Lantern” green fluorescent protein, a *RK24* oriT and the left end (LE) of the *Tn6677* transposable element. *pAra-Trans* encodes *RK24* conjugation machinery under control of an arabinose inducible promoter (P^Ara^) on an *apra^R^* (+3) single copy bacterial artificial chromosome (BAC) while the *pAra-CAST* encodes the RNA-guided *Tn6677* transposase under an arabinose inducible promoter (P^Ara^) on a *spec^R^* (+4) CDF replication backbone. The *Tn6677* crRNA is expressed under a separate constitutive promoter. Abbreviated schematics of each component is shown within paratheses next to each detailed representation. **(b**) Overview of the ACE process. The workflow starts with arbitrary cell *X* with genome *X* and cell *Y* with genome *Y* (i). *Cassette^START^* and *Cassette^END^* are sequentially integrated into genome *X* to specify the donor transfer window (ii). Engineered cell *X* is then transformed with *pAra-Trans* to be fully primed as the donor strain (iii). Cell *Y* requires no genomic modification and is transformed with *pAra-CAST* to be primed as the recipient strain (iv). For an ACE transfer experiment, donor and recipient strains are induced with arabinose and mixed in a defined ratio to allow for transfer of the oriT-flanked genomic DNA from cell *X* to cell *Y*. Transfer of genome *X* in the form of single-stranded DNA (ssDNA) initiates at the oriT in *Cassette^START^* and terminates at the oriT in *Cassette^END^* (v). Notably, transfer could also initiate at the oriT in *Cassette^END^* and terminate at the oriT in *Cassette^START^* (not depicted). The donated ssDNA recircularizes in cell *Y* using the *traI* nicking region for circularization and is converted into double-stranded DNA (dsDNA) by replication of the other strand. This circular dsDNA carries the RE and LE for recognition by the CAST complex, which integrates the dsDNA into genome *Y* in a crRNA dependent manner (vi). Plating on spectinomycin, hygromycin and sucrose enables selection for the gain and maintenance of (+2) and (+4), respectively, and the loss or absence of (−1), enabling the selection of successful post-ACE recipient cells and elimination of donor cell *X* (shown in background of vi) and virgin recipient cell without the required selection markers combination. Two integration outcomes (RL and LR integrants) in cell Y are possible due to the bidirectional transposition properties of CASTs (vii).

The donor cell is defined starting with the consecutive integrations of Cassette^START^ and Cassette^END^ to define the boundaries of conjugation (**Figure 1b ii**) and completed with the introduction of the *pAra-Trans* (**Figure 1b iii**). The recipient cell is defined with the introduction of *pAra-CAST* with a specific crRNA (**Figure 1b iv**). Upon induction of *pAra-Trans*, donor DNA is conjugated from Cassette^START^ to Cassette^END^ into the recipient cell (**Figure 1b v**). This conjugated DNA transiently circularizes in the recipient cell^36^ (**Figure 1b vi**) before it is subsequently integrated in a CAST and crRNA dependent manner, facilitated by the R-TE and L-TE sequences just within the boundaries of the transferred DNA (**Figure 1b vii**). Selection of the gain and loss of defined antibiotic markers enables selection for the desired transfer starting at Cassette^START^ and terminating at Cassette^END^. Alternative transfer starting at Cassette^END^ and terminating at Cassette^START^ is theoretically possible but is selected against and does not yield a viable intermediate for subsequent integration with the correctly positioned R-TE and L-TE. CASTs have been described to carry out bidirectional transposition, leading to two possible integration orientations, right-to-left (R-L) and left-to-right (L-R) orientations^32,33,37^ (**Figure 1b vii**).

We first validated ACE to transfer short, synthetic donor sequences up to 100 kb. To minimize any potential interference by homologous recombination, we first constructed the *E. coli* DH10B-*Syn100*Δ*lacZ* strain as the donor strain which harbors a 100-kb synthetic DNA sequence *Syn100* bearing no homology to *E. coli* genomic sequence (**Figure 2a**). ACE was accomplished with three distinct lengths of *Syn100* (10 kb, 50 kb and 100 kb) transferred from the donor to one of three target sites (TS1, TS2|TS*^lux^* and TS3) into the recipient *E. coli* DH10B (Δ*dif*::*lux*) strain (**Figure 2a-b**). TS1 and TS3 are arbitrarily chosen on opposing sides relative to the position of the origin of replication (*oriC*), while TS2|TS*^lux^* (referred from now on as TS2) is situated to target a *luxABCDE* luminesence operon replacing the original *dif* site (Δ*dif*::*lux*) in the terminus region of the chromosome. Integrations at TS2 leads to loss of chemiluminescence in this recipient strain as it knocks out the *luxA* gene in the *lux* operon.

**Figure 2.**
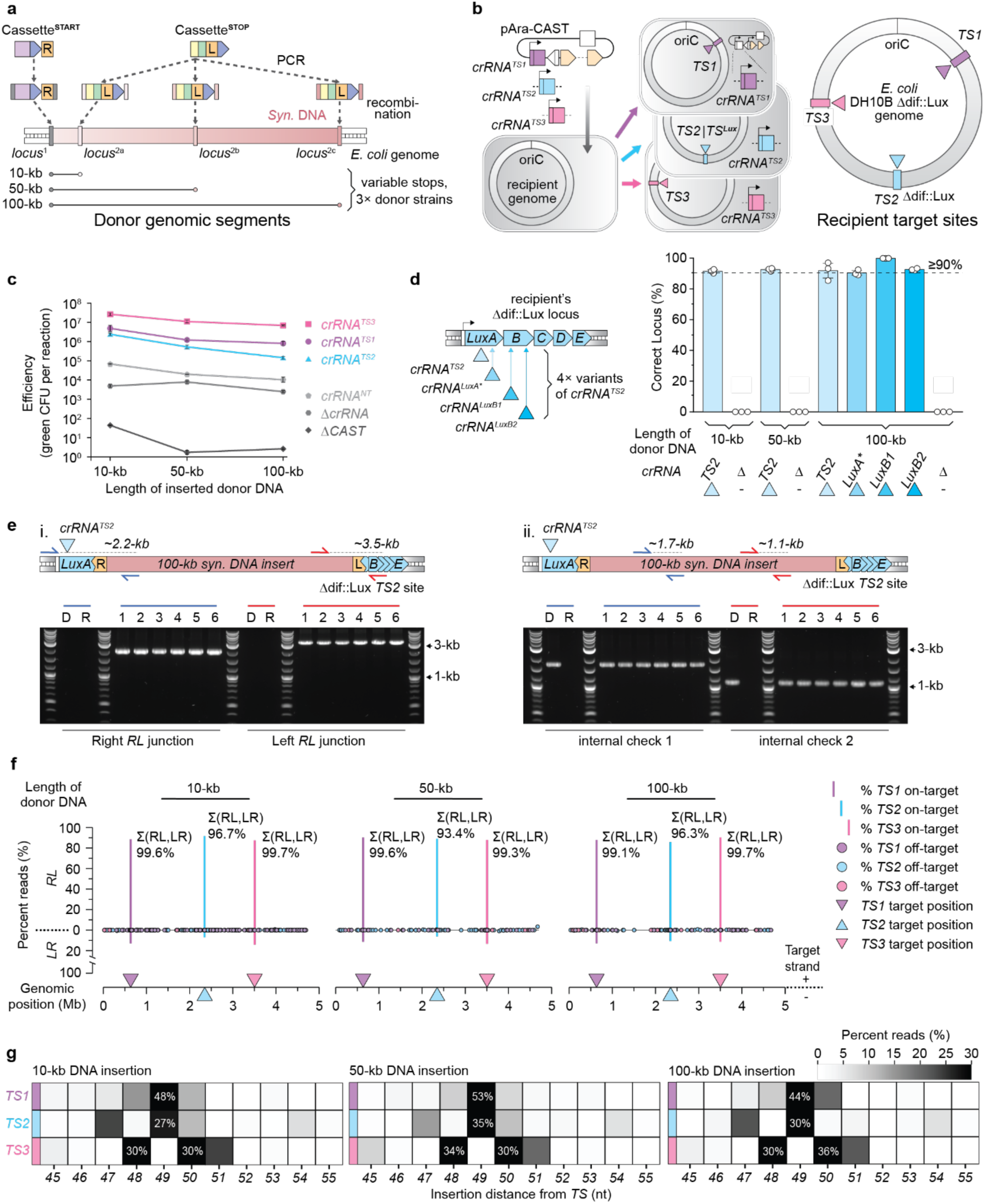
Validation and characterization of ACE. (**a**) Locations of *Cassette^START^* and *Cassette^END^*within the synthetic DNA segment *Syn100* on the genome of *E. coli* DH10B *Syn100*. *Cassette^START^* is amplified via PCR with homology arms for recombination into *locus*^1^. The location of *Cassette^END^* is variable, and is PCR amplified with the appropriate homology arms for recombination into *locus*^2a^, *locus*^2b^ and *locus*^2c^. Engineering *Cassette^STOP^* in either *locus*^2a^, *locus*^2b^ or *locus*^2c^ into *E. coli* DH10B *Syn100* with a fixed *Cassette^START^* at *locus*^1^ produces three distinct donor strains for three different transfer lengths ranging from 10 kb to 100 kb. **(b**) Preparation of *E. coli* DH10B *Δdif::lux* as recipients for integration at three defined target sites. *pAra-CAST* plasmids encoding either crRNA^TS1^ (purple square as DNA sequence on *pAra-CAST* and purple triangle as transcribed crRNA), crRNA^TS2^ (blue) or crRNA^TS3^ (pink) are transformed into *E. coli* DH10B *Δdif::lux*, generating three distinct recipients. The locations of each integration site on the *E. coli* DH10B *Δdif::lux* genome are represented by rectangles on the genome and triangles as targeting crRNA of the appropriate color. **(c**) ACE efficiency, measured in GFP+ c.f.u. per ACE experiment, of 10, 50 and 100-kb *Syn100* donors. Efficiencies were determined for transfers into recipients with crRNA^TS1^ targeting TS1 (purple line), crRNA^TS2^ targeting TS2 (blue line) and crRNA^TS3^ targeting TS3 (pink line) in addition to *pAra-CAST* variants encoding a non-targeting crRNA^NT^ (light gray line), no crRNA (ΔcrRNA, gray line), and no CAST components (ΔCAST, dark gray line). **(d**) ACE specificity at the TS2 target site for 10, 50 and 100-kb *Syn100* donors. On the left, representation of the recipient’s *Δdif*::*lux* TS2 locus showing that in addition to crRNA*^TS2^* targeting *luxA*, three other variant crRNAs targeting different portions of the operon in *luxA* and in *luxB (*crRNA*^luxA*^*, crRNA*^luxB1^*, crRNA*^luxB2^*) were used to assess transfer fidelity of the 100-kb *Syn100* donor segment. Precision of 10 and 50-kb transfers were assessed only with crRNA*^TS2^*and ΔcrRNA while 100-kb was also assessed with the three variant crRNAs. Transposition precision is calculated as the percentage of *lux*-GFP+ c.f.u. out of total GFP+ c.f.u. Data are shown as the mean ± s.d. for *n* = 3 biologically independent samples with the value of each sample independently reported as the white circle. **(e**) Genotyping colony PCR gels for six ACE integrant colonies of the 100-kb *Syn100* transfer into the *Δdif*::*lux* TS2 site. In the schematic above each gel, the red rectangle represents the 100-kb *Syn100* insert, the yellow boxes represent the LE and RE, and the blue squares rectangles and triangles represents the *lux* operon. The integration site in *luxA* targeted by crRNA*^TS2^* is shown with a blue triangle. In (i), newly formed transposition junctions in the RL integration orientation are assayed. The right and left junctions are checked with primers specific to *lux* and the *Syn100* segment, shown as blue and red arrows, respectively. In (ii), internal regions of the *Syn100* segment are assayed with two sets of primers denoted by blue and red arrows priming within *Syn100*. Bands corresponding to a set of primers are color coded accordingly. D and R denote donor and recipient controls in each assay. Lengths of each PCR product are shown above each set of arrows in the schematic. **(f**) Genome-wide distribution of Tn-seq reads, measured in percent reads, from ACE transfer experiments of 10, 50 and 100-kb cargoes into TS1 (purple), TS2 (blue) and TS3 (pink). Locations of each target site are given by triangles of the appropriate color, with the orientation of the triangle depicting the strand targeted by the crRNA. On-target transposition in the RL orientation at each site is represented by bars of the appropriate color; on-target transposition in the LR orientation are represented by bars stacked in the opposite direction. Each set of stacked bars is labeled with an on-target percentage for each target site. Off-target events, which intrinsically have an undefined integration orientation, are represented by colored circles associating them with a discrete transfer experiment. Genome coordinates are plotted with the origin of replication (*oriC*) as the zeroth coordinate. **(g**) Heat maps of integration site distributions (x-axis), measured in percent reads mapped, for 10, 50 and 100-kb *Syn100* donor cargo integrations at TS1 (purple), TS2 (blue) and TS3 (pink) as determined by Tn-seq of transfer populations.

ACE transfers of the *Syn100* are stable, specific, and efficient. With the presence of corresponding crRNAs (*crRNA^TS1^*, *crRNA^TS2^*, *crRNA^TS3^*), integration efficiencies at all target sites are high, ranging from 10^6^ to 10^8^ GFP+ c.f.u. per experiment for all sizes transferred (**Figure 2c**). With non-targeting crRNA (*crRNA^NT^*) or absence of the crRNA (*ΔcrRNA*), the c.f.u. is dramatically reduced by several magnitudes to 10^3^ to 10^5^, and further reduced to almost zero in the absence of CAST components (*ΔCAST*) (**Figure 2c**). To demonstrate the precision of ACE, we performed an *in vivo* screen leveraging concomitant loss of recipient chemiluminescence with gain of donor cargo fluorescence when integrating into the Δ*dif*::*lux* at TS2 to assay for on-target transposition (**Figure 2d, Extended Figure 1a**). For all transferred lengths, we observed a >90% ACE on-target rate consistent with previous assessments of *Tn6677* CAST specificity at shorter transfer size less than 10 kb^32,33^. This assay was repeated with three additional *lux*-targeting crRNAs (*crRNA^luxA*^, crRNA^luxB1^* and *crRNA^luxB2^*) with the 100-kb donor to control for variations caused by individual crRNA specificity (**Figure 2d**). All measured on-target rates are consistently high. We further verified the success of ACE events and the stable presence of the imported *Syn100* DNA at the correct target site by genotyping PCR of the junctions and multiple internally distributed segments across the length of *Syn100*. (**Figure 2e and Extended Figure 1b-d**). The junction PCRs also uncovered a R-L integration bias for these donors for all target locations in agreement with previous characterizations of *Tn6677* CAST system at shorter transfer size (**Figure 2e and Extended Figure 1b-d, 1f**)^32,33^.

To both quantify the observed R-L/L-R bias and improve the resolution on the on-target ACE results, we developed a tagmentation-based transposon-sequencing (Tn-Seq) protocol for genome-wide interrogation of all on and off-target ACE events (**Extended Figure 2a**). Tn-Seq of *Syn100* 10-kb, 50-kb and 100-kb transfers into TS1, TS2 and TS3 reveal on-target rate ranging from 93.4% to 99.7%, consistent with our *in vivo lux* screen on TS2 (**Figure 2f**, **Figure 2d and Extended Figure 2b and 2d**). The data also corroborates the significant R-L integration bias for all lengths transferred and locations targeted (**Figure 2f and Extended Figure 2c**). Each target had an integration distribution around the expected 49 bp insertion distance that remained constant over all tested donor sizes (**Figure 2g**)^32,33^.

ACE efficiency and specificity necessitates the presence of both crRNA and CAST components. The ability for unbiased detection of transposition throughout the genome afforded by Tn-Seq motivated us to attempt *Syn100* transfers with CAST variants containing deletions of key modules. We observed around a 10-fold decrease in efficiency for *Syn100* transfers using CAST-*crRNA^NT^*, CAST-*ΔcrRNA* and CAST-*Δcascade* and nearly a 100-fold decrease in efficiency for transfers using CAST-Δ*Qcascade* when compared to CAST-*crRNA^TS2^* (**Extended Figure 2e**). Transfers employing CAST-Δ*tns* or ΔCAST abrogated nearly all activity, demonstrating the importance of the *tnsABC* module for integration (**Extended Figure 2e**). Tn-Seq of these transfer experiments show different patterns of genome-wide non-specific integrations (**Extended Figure 2f-i, Supplemental Note 1**).

We next demonstrated ACE on a 200-kb natural segment from the partially minimized *E. coli* MDS42 genome containing a functional *lacZ* into the TS1 target region of a DH10B *lacZ^-^* recipient strain^38^ (**Extended Figure 3a**). This 200-kb donor region from MDS42 has identical sequences segregated in the recipient DH10B genome. The post ACE clones demonstrated a gain of *lacZ* activity upon plating on IPTG and X-gal (**Extended Figure 3c**). The integration success and stable maintenance of the entire 200-kb repetitive sequence was confirmed by growth phenotyping, genotyping PCR, and in particular doubled read-depth spanning duplicated sequences in Illumina sequencing (**Extended Figure 3b-f**). With this gain of 200-kb redundant sequence, we show that extensive homologies between the donor segment and recipient genome are tolerated for large ACE transfers.

### Demonstrating ACE at mega-base scale and the importance of genome architecture

The success of ACE motivated us to further scale up donor size to megabase-scale. The same *E. coli* DH10B (Δ*dif*::*lux*) strain was used as the recipient with *crRNA^TS1^*, *crRNA^TS2^*, or *crRNA^TS3^* defining the integration site at *TS1*, *TS2*, or *TS3*. The synthetic recoded *E. coli* Syn61 was chosen as the donor strain for its ∼18,000 SNPs in the form of recoding and refactoring of genes throughout its genome^14^, enabling us to reliably differentiate between the incoming donor segment and the recipient genome upon whole genome sequencing. We constructed our Syn61 donors for transfer of 100-kb, 600-kb and 1-Mb genomic DNA with Cassette^START^ and Cassette^END^ spaced roughly equidistant from the terminus region (**Figure 3a**). Three different donor sequences together with three different integration sites into the recipient genomes would yield nine possible chimeric genomes (**Figure 3b**). We observed 10^5^-10^7^ GFP+ c.f.u. for transfers of the 100-kb donor segment into various parts of the genome. The efficiencies dropped to 10^2^-10^3^ GFP+ c.f.u. for transfers of the 600-kb and 1-Mb donor segments (**Figure 3c**). As with the Syn100 donor transfers, *crRNA^NT^* or *ΔcrRNA* controls showed a 10^2^ to 10^3^-fold decrease in GFP+ c.f.u., suggesting a low off-target rate. At all lengths, *ΔCAST* controls produced close to zero GFP+ c.f.u. (**Figure 3c**). Observed genome-wide on-target rates were consistently high for ACE at megabase scale as measured by both *in vivo* chemiluminescence screen at TS2 (**Figure 3d**) and Tn-Seq at all three target positions (**Figure 3e**).

**Figure 3.**
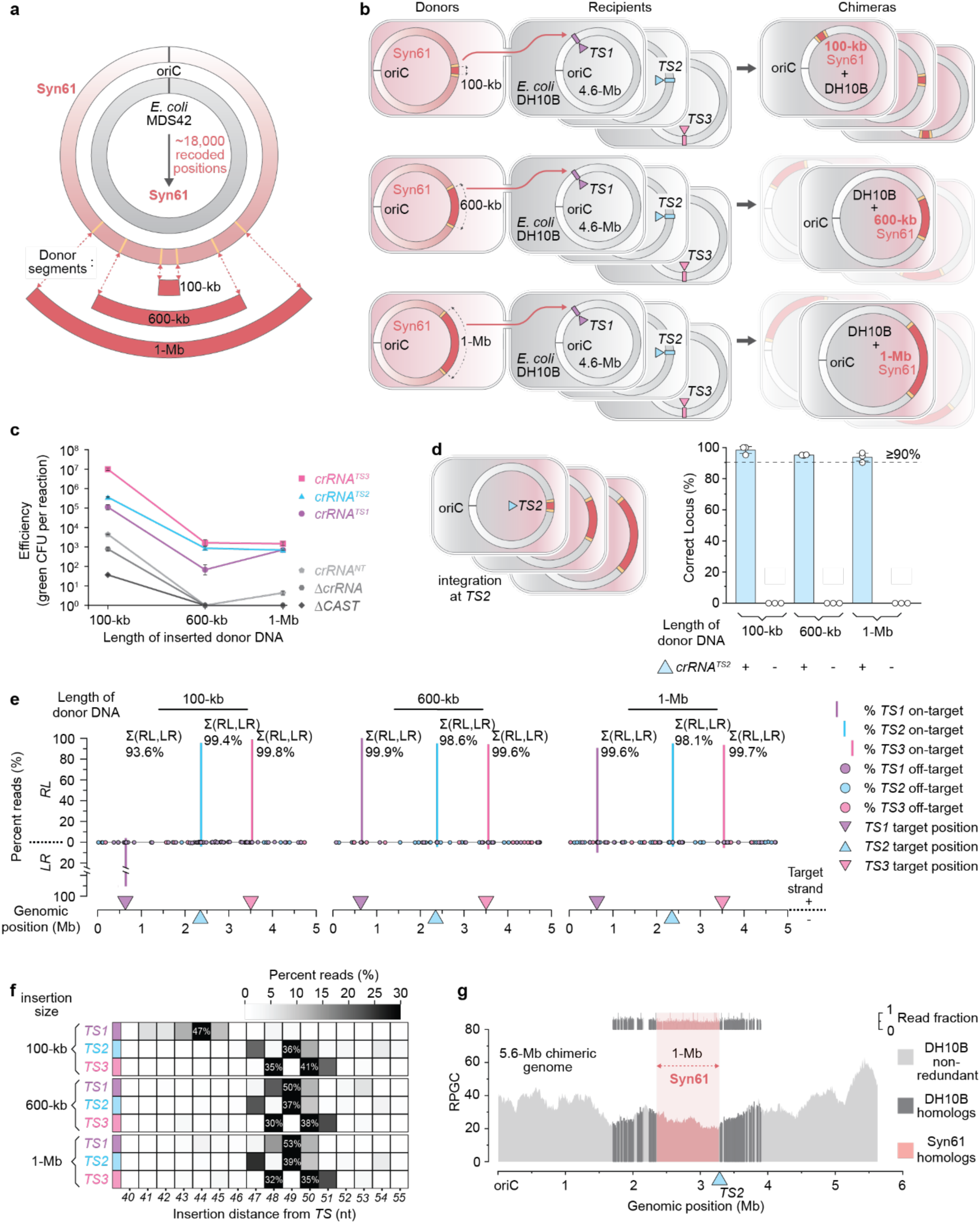
ACE enables megabase-scale additions between synthetic and natural strains. (**a**) ACE donor strain defined from the synthetic *E. coli* Syn61 strain. The recoding process converting the genome of *E. coli* MDS42 (gray ring) to *E. coli* Syn61 (pink ring) recodes approximately 18,000 codons to synonymous codons. Boundaries of the three genomic windows defined for 100-kb, 600-kb and 1-Mb transfers (red segments) centered around the *dif* site of Syn61 are defined by three sets of yellow lines on the genome. *Cassette^START^* is inserted to the yellow line right of the *dif* site and *Cassette^END^* is inserted to the yellow line left of the *dif* site. **(b**) Diagrams representing the transfer of 100-kb, 600-kb and 1-Mb Syn61 donor segments for insertion at TS1, TS2 and TS3 in recipient *E. coli* DH10B *Δdif*::*lux* strains leading to nine possible chimeric genomes. Synthesized genomes that are subsequently determined to be unstable are grayed out and set behind their stable counterparts for 600-kb and 1-Mb Syn61 transfers. Note that integration orientation of the cargo is not explicitly depicted in the final product; each chimeric genome can adopt either a RL or LR configuration. **(c**) ACE transfer efficiency, measured in GFP+ c.f.u. per ACE experiment, of 100-kb, 600-kb and 1-Mb Syn61 donors. Efficiencies were determined for transfers into recipients with corresponding crRNAs targeting TS1 (purple line), TS2 (blue line) and TS3 (pink line) in addition to *pAra-CAST* variants encoding a non-targeting crRNA^NT^ (light gray line), ΔcrRNA (gray line), and ΔCAST (dark gray line). **(d**) ACE specificity at the TS2 target site for 100-kb, 600-kb and 1-Mb Syn61 donors. On the left, representation of the recipient *Δdif*::*lux* TS2 locus with 100-kb, 600-kb and 1-Mb Syn61 cargoes (in red) integrations. Unlike in **2d**, Precision of megabase-scale cargo integration was only assessed with crRNA*^TS2^*and ΔcrRNA. Transposition precision is calculated as the percentage of *lux*-GFP+ c.f.u. out of total GFP+ c.f.u. Data are shown as the mean ± s.d. for *n* = 3 biologically independent samples with the value of each sample independently reported as the white circle. **(e**) Genome-wide distribution of Tn-seq reads, measured in precent reads, from ACE transfer experiments of 100-kb, 600-kb and 1-Mb donor cargoes into TS1 (purple), TS2 (blue) and TS3 (pink). On-target transposition, off-target events, transposition orientation, crRNA target locations and genome coordinates are represented and plotted as in **2f**. **(f**) Heat maps of integration site distributions (x-axis), measured in percent reads mapped, for 100-kb, 600-kb and 1-Mb donor cargo integrations at TS1 (purple), TS2 (blue) and TS3 (pink) as determined by Tn-seq of transfer populations. **(g**) Genome-wide read coverages (in RPGC) for the *E. coli* DH10B + 1-Mb Syn61 genomic chimeric strain. Syn61 genomic segments are shaded in pink, and homologous *E. coli* DH10B genomic segments are shaded in dark gray. Non-redundant sections of the *E. coli* genome are in lighter grays. Location of the TS2 target site is given by the blue triangle. Read fraction mapping to the Syn61 segment and the homologous DH10B segment is set above the coverage plot. Read fractions share the same color coding as the coverage plot.

We observed R-L integration bias again for all lengths and locations transferred, except in the unique exception of 100-kb donor segment into recipient genome position TS1, which had a predominantly flipped L-R integration bias (**Figure 3e**). Examination of the 100-kb donor segment reveals the presence of a single *ter* site *terC*. *Ter* sequences situate throughout the terminus region of the *E. coli* genome and have been implicated in the termination of genome replication by forming unidirectional replication fork traps when bound by the protein *tus*^39,40,41,42^ (**Extended Figure 4a**). Consistent with the required orientation of the *ter* sites, we realized that L-R integration of the 100-kb donor in TS1 orients the *terC* in that segment in alignment with the required DNA replication directionality, suggesting that R-L integration would lead to replication fork arrest at the non-permissive end of the inverted *terC* (**Extended Figure 4b**). This deviation from R-L integration bias for this condition was also accompanied by a different integration distribution centered around 44 bp (**Figure 3f**). Otherwise, all other conditions had expected distributions in agreement with previous *Syn100* measurements (**Figure 3f**, in comparison to **Figure 2g**).

Successful ACE events are necessary but may not be sufficient for long-term stability of the fused chimeric genome. We verified the stability of post-ACE clones with next generation sequencing. Whole genome sequencing showcased the presence and integrity of the entire transferred segment in the correct location for 100-kb, 600-kb and 1-Mb Syn61 donor segments when integrated into the recipient’s TS2 position (**Figure 3g and Extended Figure 4b, 5a-b**). The 100-kb donor segment was also stable in the recipient’s TS1 and TS3 genomic positions with the requirement of a properly oriented *terC* site. In contrast, the 600-kb and 1-Mb donor segments, centering the terminus region of the donor genome and both containing multiple *ter* sites facing opposite directions, were not maintained in the recipient genome upon integration into TS1 or TS3 positions (**Extended Figure 4b and 5a-b**). As in the case of R-L 100-kb TS1 integration, this observation is consistent with the hypothesis that integration of 600-kb and 1-Mb donor segments in either orientation presents the post-ACE chimeric chromosome with numerous opposing *ter* sites orientated towards their non-permissive directions, trapping the advance of the replisome during normal chromosome replication^39,43^ (**Extended Figure 5a-b**). We thus noted the important distinction between on-target ACE integration and stable post-ACE maintenance of the donor DNA. While the Tn-Seq data shows that the transfers are generally on-target in all conditions, we infer that a majority, if not all, of those on-target integrations of donated segments with non-permissive architectures involve the subsequent loss of the whole or part of the internal donor segments possibly due to genomic instability from premature replication fork trapping^43,44^.

We were also able to utilize the SNPs distributed in the coding sequences of the incoming Syn61 donor to differentiate its recoded genes from their wild-type DH10B homologs. Through this variant calling analysis, the identity of successfully integrated and stably maintained donor segments of 100-kb, 600-kb and 1-Mb Syn61 genomic segments at the recipient DH10B’s TS2 target was confirmed, revealing general parity between the frequency of the wild-type DH10B genes and their recoded Syn61 homologs, each at their expected locations within the chimeric genome (**Figure 3g and Extended Figure 4b, 5a-b**).

### ACE at multi-megabase scale and modularity across the entire donor genome

We next systematically explored the possibility of mobilizing DNA from all regions of the donor genome. To enable this, we engineered multiple Syn61 donors that each would mobilize a very large discrete segment of their synthetic genomes in an ACE event.

To push the limits of ACE, the first donor segment (seg. 1) spans 2-Mb, or half of its genome, centering around the *dif* terminus region (**Figure 4a**). Donor seg. 2 covers 1-Mb segment downstream from the *oriC*, and seg. 3+4 covers 1-Mb upstream of *oriC* (**Figure 4a**). An additional 1-Mb donor segment seg. 5 is defined spanning across *oriC* (**Figure 4a**). Segments seg. 2, seg. 3+4 and seg. 5 do not contain any primary *ter* sequences. Seg. 3+4 is further divided into two 500-kb segments as separate seg. 3 and seg. 4. Together, these donor segments span the entirety of the 4-Mb donor genome (**Figure 4a**).

**Figure 4.**
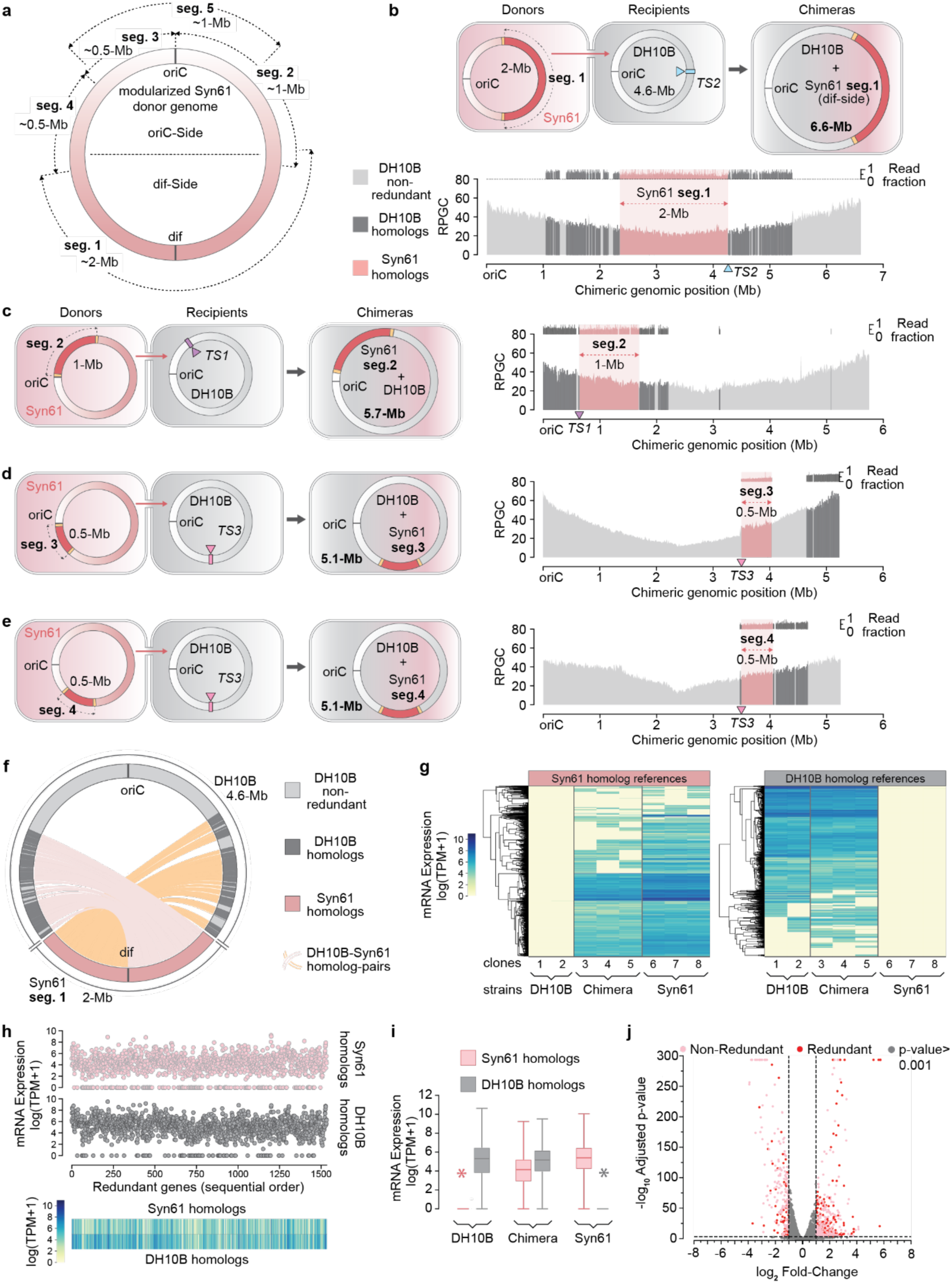
Modularity and scalability of ACE to create transcriptionally active genomic chimeras. (**a**) Division of the *E. coli* Syn61 genome into multiple synthetic segments (seg. 1 to 5) for mobilization in discrete donor strains. Seg. 1 encompasses the *dif*-side of the genome and is ∼2 Mb, while seg. 2 encompasses the right side of the *oriC* and is ∼1 Mb. Seg. 3 and 4 together encompass the left ∼ 1 Mb side of the *oriC* and are ∼500 Kb each. Seg. 5 spans across the *oriC* and is ∼1 Mb. **(b**) Top: Diagram of ∼2-Mb seg. 1 transfer from *E. coli* Syn61 to DH10B *Δdif*::*lux*. The Syn61 genome is depicted by the pink circle and the *E. coli* DH10B genome are depicted by the gray circle. Sizes (in Mb) of transferred region, recipient genome and chimeric genome are given in the schematic. The transferred region is centered around the *dif* site of Syn61 and is represented by the red portion of the circle flanked by TEs, shown as yellow rectangles. Transfer direction is shown by the red arrow. The TS2 site of *E. coli* DH10B *Δdif*::*lux* is given by the blue rectangle, which is targeted by the CAST by crRNA^TS2^ shown as a blue triangle. The outcome chimeric genome (6.6 Mb) is represented as an integration of the red segment of the *Syn61* genome into the gray *DH10B* genome at the *Δdif*::*lux* site. Bottom: Genome-wide read coverages (in RPGC) for the *E. coli* DH10B *Δdif*::*lux* + 2-Mb seg. 1 genomic chimeric strain. Syn61 genomic segments are shaded in pink, and homologous DH10B genomic segments are shaded in dark gray. Non-redundant sections of the *E. coli* genome are in lighter grays. Location of the TS2 target is given by the blue triangle. Read fraction mapping to seg. 1 and the homologous DH10B segment is set above the coverage plot. Read fractions share the same color coding as the coverage plot. **(c**) As in **b**. but transfer diagrams of 1-Mb seg. 2 donor for insertion at TS1 (purple) in recipient *E. coli* DH10B *Δdif*::*lux* strains (left) and associated genome-wide read coverage plot and read fraction mapping for the resultant chimeric genome (right). **(d**) As in **b**. but transfer diagrams of 500-kb seg. 3 donor for insertion at TS3 (pink) in recipient *E. coli* DH10B *Δdif*::*lux* strains (left) and associated genome-wide read coverage plot and read fraction mapping for the resultant chimeric genome (right). **(e**) As in **b**. but transfer diagrams of 500-kb seg. 4 donor for insertion at TS3 (pink) in recipient *E. coli* DH10B *Δdif*::*lux* strains (left) and associated genome-wide read coverage plot and read fraction mapping for the resultant chimeric genome (right). **(f**) Genome-wide mapping of Syn61 gene variants given in pink to their corresponding DH10B wild-type homologs given in dark gray in the *E. coli* DH10B *Δdif*::*lux* + 2 Mb seg. 1 chimeric genome with a R-L integration orientation. Pairs of DH10B and Syn61 homologs are connected by either pink or yellow lines. **(g**) Hierarchical clustering of transcript counts, measured in log of transcripts per million + 1 (log(TPM+1)), of Syn61 gene homologs and DH10B gene homologs in the DH10B, Chimeric and Syn61 strains. Up to three individual replicates are shown for each synthetic strain. Genes are clustered based on expression level within each genomic background. **(h**) Genome-wide scatterplot (top) and heat map (bottom) of positional expression, in log(TPM+1), of each Syn61 gene homolog in pink and DH10B gene homolog in dark gray within the chimeric DH10B + 2 Mb Syn61 strain. Genes are plotted in positional order as they appear on the genome. **(i**) Expression distribution of bulk Syn61 transcripts (pink) compared to bulk homologous DH10B transcripts (dark gray) in DH10B, Chimeric and Syn61 strains measured in log(TPM+1). Distributions are based on the data from *n = 3* biologically independent samples for the Chimera and Syn61 strains and *n = 2* biologically independent samples for the DH10B strain. Asterisks represent outliers that are the result of bioinformatic and sequencing artifacts with no biological significance. **(j**) Differential expression of redundant genes (red) and non-redundant genes (pink) between the chimeric strain and the DH10B background strain. The log_2_ of the ratio of the expression level of the redundant genes in the chimeric strain to the expression level of their homolog in the DH10B strain (red) is plotted alongside log_2_ of the ratio of the expression level of non-redundant genes in the chimeric strain to the expression level of its homolog in the DH10B strain (pink). Points are also plotted based on their – log_10_ adjusted p-value. P-value cutoff was set at 10^-3^ and the log_2_ fold-change cutoff was set to ±1 so that points below the cutoff are in gray.

**Figure 5.**
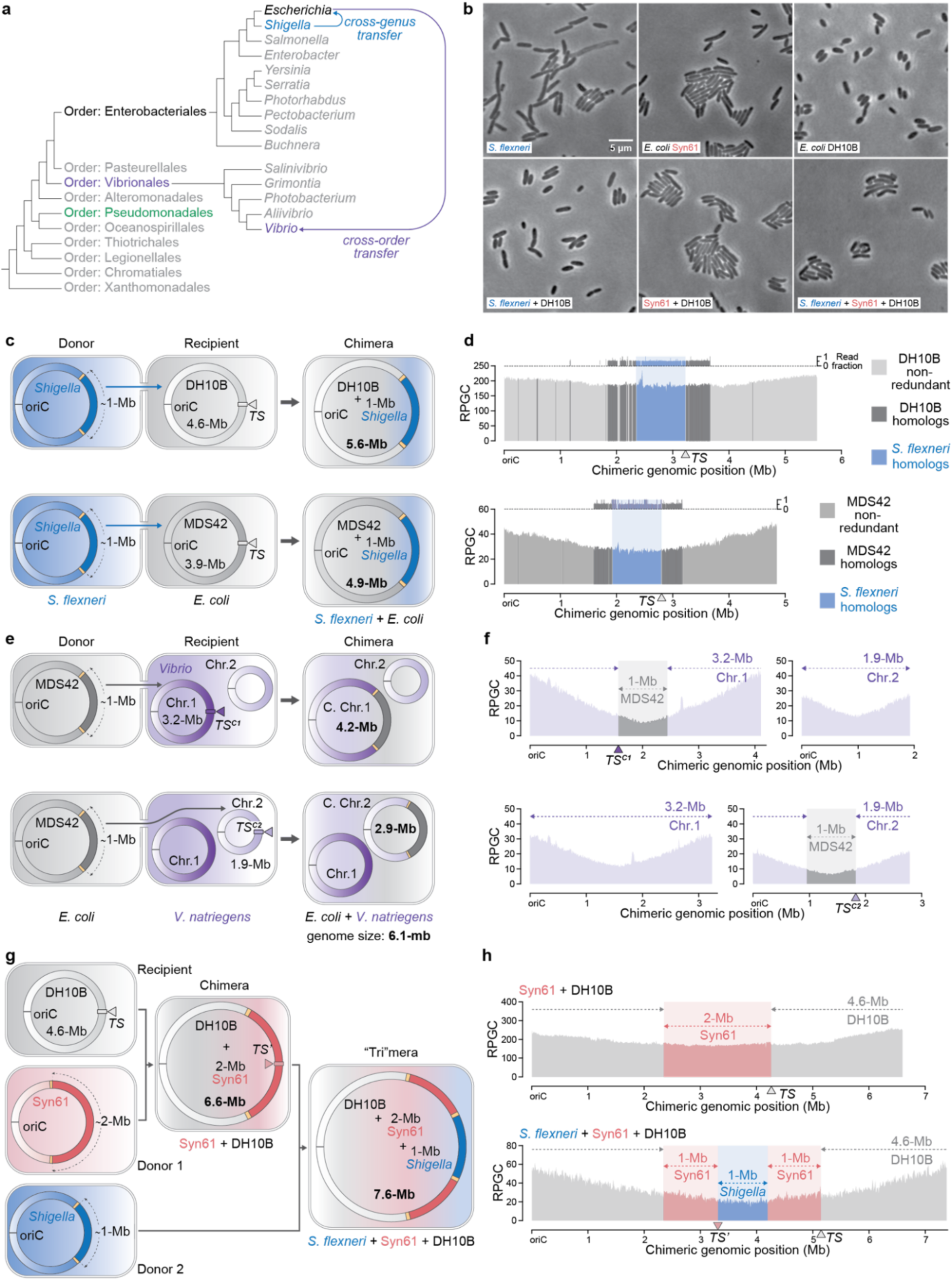
Creation of cross-species/genus/order genomic chimeras and trimeras. (**a**) Phylogenetic tree within the class of *Gammaproteobacteria*, specifically between *Escherichia* and *Shigella* genera and between the *Enterobacteriales* and *Vibrionales* orders. Cross-genus DNA transfer from *Shigella* to *Escherichia* is shown by the blue arrow. Cross-order DNA transfer from *Escherichia* to *Vibrio* is depicted by the purple arrow. The *Pseudomonadales* order is also bolded with green to emphasize its successful use as a recipient in an ACE experiment. **(b**) Phase contrast confocal microscopy of synthesized cross-strain and cross-species genome chimeras. Scale bar is provided in the first image of *S. flexneri*. Strains and chimeras are color-coded according to the following: blue for *S. flexneri*, pink for Syn61 and black for *E. coli* DH10B. **(c**) Diagram of ∼1-Mb *S. flexneri* donor fragment transfer to *E. coli* DH10B and *E. coli* MDS42. The *S. flexneri* genome is depicted by the blue circle and the *E. coli* genomes are depicted by the gray circle. Sizes (in Mb) of transferred region, recipient genome and chimeric genome are given in the schematic. The transferred region is centered around the *dif* site of *S. flexneri* and is represented by the dark blue portion of the circle flanked by TEs, shown as yellow rectangles. Transfer direction is shown by the blue arrow. The *dif* site of *E. coli* is given by the gray rectangle, which is targeted by the CAST by the TS crRNA, shown as a gray triangle. The outcome chimeric genome (5.6 Mb for DH10B*-S. flexneri* and 4.9 Mb for MDS42*-S. flexneri*) is represented as an integration of the dark blue segment of the *S. flexneri* genome into the gray *E. coli* genome at the *dif* site. **(d**) Genome-wide read coverages (in RPGC) for the *E. coli* DH10B + 1-Mb *S. flexneri* (top) and *E coli* MDS42 + 1-Mb *S. flexneri* (bottom) genomic chimeric strain. *S. flexneri* genomic segments are shaded in blue, and homologous *E. coli* genomic segments are shaded in dark gray. Non-redundant sections of the *E. coli* genome are in lighter grays. Location of the TS *dif* target is given by the gray triangle. Read fraction mapping to the *S. flexneri* segment and the homologous DH10B segment is set above the coverage plot. Read fractions share the same color coding as the coverage plot. **(e**) Diagram of 1-Mb *E. coli* MDS42 donor fragment transfer to *V. natriegens*. The MDS42 genome is depicted by the gray circle and the *V. natriegens* genomes are depicted by the purple circles, with chromosome 1 in dark purple and chromosome 2 in light purple. Sizes (in Mb) of transferred region, recipient genome and chimeric genome are given in the schematic. The transferred region is centered around the *dif* site of MDS42 and is represented by the dark gray portion of the circle flanked by TEs, shown as yellow rectangles. Transfer direction is shown by the gray arrow. The *dif* site of *V. natriegens* chromosome 1 is given by the dark purple rectangle, which is targeted by the CAST by the TS^C1^ crRNA, shown as a purple triangle. The *dif* site of *V. natriegens* chromosome 2 is likewise represented by a purple rectangle targeted by TS^C2^ shown as a purple triangle. The outcome chimeric genomes (6.1 Mb) are represented as an integration of the dark gray segment of the MDS42 genome into either chromosome 1 or chromosome 2 of the *V. natriegens* genome at their respective *dif* sites. **(f**) Genome-wide read coverage (in RPGC) for the *V. natriegens* chr.1 *+* 1-Mb MDS42 and *V. natriegens* chr.2 + 1-Mb MDS42 chimeric genome strain. MDS42 genomic segments are shaded in gray, *V. natriegens* genomic segments are shaded in purple. The sizes and spans of each section of the chimeras are given by dotted lines of the appropriate color. The genomic locations of TS^C1^ and TS^C2^ are given by dark purple and purple triangles. **(g**) Diagram of the stepwise ACE synthesis of a tripartite genomic hybrid (trimera) between *E. coli* DH10B, *E. coli* Syn61 and *S. flexneri*. Sizes (in Mb) of transferred region, recipient genomes and chimeric/trimeric genomes are given in the schematic. In the first step, the creation of *E. coli* DH10B + 2-Mb Syn61 seg. 1 is as previously described in **4b**. with the exception of the recipient in this case being the wild-type DH10B without *Δdif::lux* with a TS crRNA. The ∼1-Mb *S. flexneri* donor fragment, as previously seen in **c**. is then integrated in the second step into *E. coli* DH10B + 2-Mb Syn61 at the *dif* site of the chimera as depicted with the pink rectangle with the TS’ crRNA, represented by a pink triangle. The outcome trimeric genome (7.4 Mb for DH10B*-*Syn61*-S. flexneri*) is represented as an integration of the dark blue segment of the *S. flexneri* genome into the DH10B *+* 2 Mb Syn61 genome at the *dif* site. **(h**) As in **d**. but the genome-wide read coverage (in RPGC) for the *E. coli* DH10B *+* 2-Mb Syn61 chimeric strain and *E. coli* DH10B + 2 Mb Syn61 + 1 Mb *S. flexneri* trimeric genome strain. *S. flexneri* genomic segments are shaded in blue, Syn61 genomic segments are shaded in pink and DH10B genomic segments are shaded in gray. The sizes and spans of each section of the trimera are given by dotted lines of the appropriate color. The genomic locations of TS and TS’ are given by gray and pink triangles.

The integration of the 2-Mb seg. 1 donor segment into the recipient’s TS2 genomic position would constitute an almost 50% expansion of the *E. coli* genome. Amazingly, this chimeric genome was characterized to be successful and stable via whole genome sequencing, (**Figure 4b and Extended Figure 7d-f**). Efforts to integrate the same 2-Mb seg. 1 into the TS1 and TS3 sites failed to yield stable clones (**Extended Figure 7g-h**), possibly due to the same forbidden genomic architecture with opposing directions of *ter* sites previously described.

We tested the transfer of each distinct donor segment into recipient DH10B Δ*dif*::*lux* genome at TS1, TS2 or TS3 positions (**Figure 4c, d, e and Extended Figure 7a**). While the 1-Mb seg. 2 can be stably integrated into all three sites in the recipient’s genome, the seg. 3+4 of the same size failed to generate viable clones. (**Figure 4c, Extended Figure 6a-b and data not shown**). After splitting the single 1-Mb contiguous seg. 3+4 segment into the two separate 500-kb segments seg. 3 and seg. 4, integrations of both seg. 3 and seg. 4 yielded stable post-ACE clones into all three sites in the recipient’s genome (**Figure d, e and Extended Figure 6a, 6c-d**). While it has been suggested that maintaining symmetry between the replichores is important for bacterial genome stability^29,44,45^, our observations suggest that the bacterial genome is robust to at least tolerate large megabase-scale insertions at various positions on the chromosome, provided the absence of misaligned *ter* sites relative to the direction of the replichore.

Additionally, we report the successful transfer of the 1-Mb seg. 5 centered around the *oriC* of Syn61 into TS3 of a DH10B recipient. This procedure generated genome chimeras that have an additional, ectopic replication origin. Whole genome sequencing indicates that this inserted secondary *oriC* is active, due the presence of a maxima of read coverage at this *oriC* (**Extended Figure 7a-c**).

Collectively taken, we show that the entirety of a bacterial genome can be transferred in discrete sections into other bacteria to build genome chimeras.

### Transcriptomics of chimeric genomes fused by ACE

The creation of a 4.6-Mb *E. coli* DH10B + 2-Mb Syn61 genome chimera presents a unique opportunity to investigate its transcriptomic profile. This first-in-class artificial bacterial strain has nearly half of its gene content complemented by recoded homologs (**Figure 4f**). This will allow us to understand whether these recoded homologs express and how their presence influences native genomic context.

RNA-Seq measurements were performed on said chimeric genome alongside donor and recipient controls. An initial inspection of the data suggests that imported recoded complements and their native homologs are both transcriptionally active when comparing individual replicates through hierarchical clustering and grouped datasets through positional expression (**Figure 4g and 4h**). A comparison of the expression distribution for both the DH10B genes and Syn61 genes uncovers no significant difference between the levels of expression for Syn61 and DH10B gene homolog categories (**Figure 4i**). Differential expression of the chimeric genome compared to the DH10B background reveals an overexpression bias of at least 1-fold change for a subset of genes that were duplicated and made redundant via ACE (**Figure 4j**). Nonetheless, the strength of expression changes observed in our data are not solely determined by the copy number of genes, and deeper exploration is required to understand the regulatory dynamics of chimeric genomes constructed via ACE. We can consequently conclude that the massive complementation of the DH10B genome with recoded genes from Syn61 leads to expression of the introduced genes and perturbation of the native genomic background.

### Cross-species megabase genomic additions

We went on to investigate whether ACE was generalizable to donor and recipient bacteria beyond *E. coli* strains for the creation of cross-genus genome chimeras. Towards this aim, we engineered an attenuated^46,47^ *Shigella flexneri* strain (*S. flexneri* CFS100) into a donor to donate about 1 Mb of its genome^48^. *S. flexneri* was chosen as *Shigella* is classified to be the genus closest related to the genus of *E. coli* (**Figure 5a**). In a single ACE step, the engineered *S. flexneri* donor strain successfully transferred 900-kb of its genome centered around the *dif* into the *dif* site of an *E. coli* DH10B recipient (**Figure 5c-d**). The success and stability of this cross-genus chimeric genome was validated by whole genome sequencing (**Figure 5d**). A corresponding transfer of the *S. flexneri* donor using a different *E. coli* strain MDS42 *dif* was also performed and confirmed to have been successful (**Figure 5c-d**).

RNA expression profiles of both the DH10B-*Shigella* and the MDS42-*Shigella* ACE chimeras were interrogated in this work (**Extended Figure 8a-e**). Similar to our observations for the DH10B-Syn61 ACE chimera, DH10B– and MDS42-*Shigella* ACE chimeras showcase robust transcriptional expression for both imported gene homologs and their native complements (**Extended Figure 8b and 8c**). Novel *Shigella* genes that were gained through ACE expressed readily in both DH10B and MDS42 recipient backgrounds (**Extended Figure 8d**). Expression of genes originally deleted in the MDS42 strain background were reconstituted upon chimerization via ACE (**Extended Figure 8e**). We additionally carried out differential expression analysis for the DH10B– and MDS42-*Shigella* ACE chimeras compared against DH10B background. This revealed a strong bias of at least 1-fold change for genes that were duplicated and made redundant via ACE (**Extended Figure 8f**). Of note, gene ontology analysis of the intersection of overexpressed genes in the DH10B– and MDS42-*Shigella* ACE chimera showcased strong upregulation of genes involved in iron-sulfur complex assembly, glyoxylate bypass, peroxidase, and flagellar assembly (**Extended Figure 8g and 8h**).

ACE was also tested across more distantly related bacteria. We tested ACE transfers into industrially relevant bacteria *Vibrio natriegens* and *Pseudomonas putida*^49,50^. Compared to *S. flexneri*, these strains are more phylogenetically distant to *E. coli* and are classified in distinct orders (**Figure 5a**). We initially confirmed transfers of the 200-kb MDS42 donor segment into a previously validated insertion site of the primary chromosome of the *Vibrio* genome^51^ (**Extended Figure 9a-d**). Excitingly, transfers of a 1-Mb MDS42 donor segment centered around the *dif* were achieved in both the *dif* site of chromosome 1 and of chromosome 2 in *V. natriegens* (**Figure 5e-f and Extended Figure 9e-h**). However, although being viable and stable from room temperature to 37°C, these chimeric strains suffered decreased fitness at 4°C and were not culturable after long term cryogenic storage, consistent with previous reports of the susceptibility *V. natriegens* to cold temperatures^49^. Pilot transfers of 100-kb *Syn100* were also successful from *E. coli* donor into *P. putida* recipient when targeting downstream of the *glmS* gene (**Extended Figure 9i-k**). In summary, we demonstrate proof of concept ACE transfers of large donors across diverse bacteria spanning different species, genus, and orders, providing first-in-class evidence towards the functional expression of megabase-scale cross-species genomic hybrids.

### Iteration of megabase transfers: Creation of a genomic trimera

We further demonstrated the iteration of ACE to enable even larger scale sequential additions. The ability to iteratively transfer multiple megabases from different donor bacteria would significantly expand the number and complexity of artificial genome chimeras attainable through this method. We therefore asked whether multiple large insertions with ACE are possible on the same recipient.

Towards this end, we decided to test the iterative integration of the 900-kb *S. flexneri* donor segment into the previously validated *E. coli* DH10B + 2-Mb Syn61 seg. 1 chimeric strain (**Figure 5g**). Anticipating this, we had previously flanked the +2/-2 GFP selection/screening cassette adjacent to the L-TE in the Cassette^END^ with recombinase sites of the *PhiC31* integrase^52^. Expression of the integrase successfully excised the +2/-2 markers in addition to the *GFP* from the genome of *E. coli* DH10B + 2-Mb Syn61 seg. 1, priming this chimeric strain for subsequent megabase-scale ACE transfer (**Extended Figure 10a-b**). We reasoned that we would not encounter the phenomenon of Tn7-like transposon target immunity if we integrate the *S. flexneri* donor into the middle of the 2-Mb Syn61 segment of the chimera using the *dif* crRNA^33,53^. The R-TE and L-TE of the newly integrated donor would be approximately 1 Mb from both the R-TE and L-TE of the previous integration, thus avoiding a reduction of transposition efficiency. The sequential integration of a 1-Mb *S. flexneri* donor segment into the *dif* site of the Syn61-DH10B chimeric genome yielded stable trimeric genomes (**Figure 5h and Extended Figure 10c-d**).

This stable trimeric genome was then interrogated in the context of RNA transcriptomics, and all three homolog gene variants were shown to be expressed robustly (**Extended Figure 10e**). Interestingly, the RNA count distribution highlights an order of preference for the strength of expression of the homolog variants, where DH10B native variants are expressed more than Syn61 homolog variants, and where Syn61 homolog variants are expressed more than *Shigella* variants. Differential expression was carried out comparing the trimeric strain to the DH10B background; our findings suggest that – alike previous ACE chimeric strains – there is a non-deterministic enrichment in gene overexpression above 1-fold for both doubly redundant and triply redundant genes (**Extended Figure 10f**).

Confocal microscopy of these chimeric and trimeric strains revealed rod-shaped morphologies akin to their wild-type bacterial progenitors (**Figure 5b**). Growth assays of these strains also reveal that larger and successive transfers slow the growth rate of these strains to a moderate extent (**Extended Figure 10g-h**). Taken together, we present the successful creation of a cross-strain cross-genus genome trimera with a remarkably expanded genome of ∼7.5 Mb, a ∼3 Mb addition of genetic information in two successive ACE steps.

## Discussion

ACE enables multi-megabase integrations across bacterial genomes to create synthetic chimeras in a single step, while iterations of ACE create trimeric strains larger than any prior reports. These GEOs generated by ACE were stable and transcriptionally active. ACE functions across strains, species, genus, and orders, illustrating the feasibility of rapid, robust, affordable and scalable construction of artificial life.

The success of ACE depends on the unique design coupling the oriT and TE sequences to specify the transfer window and facilitate the crRNA guided integration. Initiation of conjugation at the first starting oriT in Cassette^START^ and termination of conjugation at the second ending oriT in Cassette^END^ enables tight regulation of what is transferred and integrated to the recipient, and subsequently the oriT-adjacent TEs are used for targeted insertion. We are aware that this double-oriT configuration leads to two possible stretches of transferred DNA, hence the specific positions of the R-TE /L-TE and introduction of markers to counter-select against the undesired product starting from Cassette^END^ and ending at Cassette^START^. While this is effective for substrates of lengths equal to or less than half the size of the donor genome, we realize this double-oriT approach may reduce the efficiency of transfer for substrates larger than half the size of the donor genome. Engineering the conjugation machinery and the oriT sequences may pave the way to higher efficiency and greater robustness especially for greater than ½ genome ACE operations.

The ACE design is also conducive to biocontainment as it decouples conjugation of the donor genomic fragment and integration of the transferred fragment into two separate plasmids in two cellular compartments. The donor DNA does not encode the means of its propagation after the conjugation into another cell without *pAra-Trans*. Only the designated recipients that carry the *pAra-CAST* will be able to incorporate the donated DNA, otherwise the transferred DNA cannot replicate and maintain itself. Hence, ACE has been inherently structured to limit the inadvertent propagation of transferred DNA outside of its intended target.

ACE demonstrates the capacity of CAST on the multi-megabase scale. By incorporating type I-F CAST as part of our ACE system, we expanded its targeted genomic integration capacity from 10-kb all the way to 2-Mb, ∼200-fold greater than previously reported^33^. Our findings suggest that the multi-stage recruitment of the cascade-tniQ-tnsC-transposome complex in the type I-F CAST, shown to mediate licensing of accurate target site selection and transposition^54,55,56^, remains functional even in the context of large genome-scale donor segments.

ACE offers a powerful alternative to homologous recombination-based methods. Genome integration by ACE via CAST inherently bypasses the need for double strand breaks and/or homologous recombination for incorporating changes into a chromosome^33^. By building cross-strain genomes via ACE, we largely minimized genomic instability and undesired random recombination events, the occurrence of which is typically elevated when using homologous recombination and double strand breaks dependent techniques to transfer megabase-sized DNA inserts containing long stretches of redundant DNA homologous to the recipient genome.

While specific, CAST mediated transposition is also flexible in its programmable targeting. The reported promiscuity in PAM requirements of Type I-F CASTs enables use of crRNAs with non-canonical PAMs^32,56,57^. Indeed, in this study we deviate in certain occasions from the 5’-CC-3’ PAM requirement, such as using a 5’-CA-3’ PAM when targeting the *dif* site, without noticeable reductions in ACE efficiency. Portability of Type I-F CASTs, as demonstrated in human cells^58^, may also enable translation of ACE into higher eukaryotes, despite the challenge of megabase DNA cargo delivery. We are nonetheless curious of the possibilities of translating this technology into more complex systems and its implications for engineering synthetic eukaryotic genomes and therapeutics.

ACE can also be configured to use other types of CASTs. Although we chose to use a member of the Type I-F multi-subunit effector CAST family for use in ACE single-effector CASTs, such as those of the Type V-K system, could theoretically replace the Type I-F system as the integration module in this protocol^59^. We have conducted preliminary experiments using Cas12k-TniQ-TnsBC in a similar conjugative workflow using Cas12k coupled transposases to successfully transfer *Syn100* but chose to continue with the Type I-F on account of its greater integration fidelity and absence of co-integrate formation^60^ (data not shown). However, numerous advances in engineering target selection and eliminating co-integrate products in Type V-K CASTs may elevate conjugation coupled Type V-K transposition as a suitable alternative to Type I-F CAST in ACE^61, 62^.

With numerous benefits, ACE offers improved alternatives to all steps of the current genome synthesis paradigm. Taking advantage of life’s natural diversity, artificial genomes with designated properties may not need to be designed from scratch and then painstakingly synthesized and assembled, but rather melded together in a single ACE or an iteration of several ACE steps. Moreover, by leveraging engineered conjugative delivery, we can transfer multi-megabase of DNA between bacteria in a single ACE step, in contrast to the low efficiencies of *in vitro* delivery of 100-kb or larger synthetic DNA. Finally, as a novel universal way of inserting syn. DNA, ACE is the first reported use of crRNA guided CAST integration for building synthetic genomes, enabling stable addition and maintenance of multi-megabase of highly homologous genomic segments.

By encoding complex engineered functions into chimeric genomes, ACE has the potential to spawn applications where these large truly genomic scale insertions can confer novel phenotypes and physiology for therapeutic and industrial applications. For example, ACE allows for transfers of large biosynthetic gene clusters into a variety of heterologous hosts for cluster expression, characterization, and refactoring for metabolic engineering^63,64^. Beyond transferring natural DNA, ACE can also transfer synthetic DNA, interfacing with a plethora of *in vitro* and *in vivo* DNA assembly techniques. The ability to integrate large *de novo* designed and assembled DNA, combined with downstream transcriptomic, proteomic and phenotyping characterization can accelerate exploration of synthetic genome-scale modules for encoding desired functionalities^65,66^.

Large genomic transfers with ACE reveals patterns in genome architecture. Our integration experiments of segments centered around the *dif* and adjacent to the *oriC* into targets around the recipient *E. coli* genome enabled us to uncover the role of length and identity of the donor fragment in determining permissive transfer sites. We observed that donor segments centered around the *dif* and containing *ter* sites can be integrated but were unstable in target regions where the *ter* sites have their non-permissive ends orientated towards the *oriC*. We hypothesize that these ectopic *ter* sites prematurely arrest replication forks before they can reach the natural replication fork trap region of the genome opposite of the *oriC*, leading to incomplete genome replication, chromosomal instability, and subsequent rearrangement^39,41,42,43^. Moreover, donors chosen adjacent to the *oriC* lacking *ter* sequences were largely stably integrated throughout the genome, although at variable rates, but nonetheless suggest a tolerance of the *E. coli* chromosome for large replichore asymmetries^44,45^. Elucidating more precise rules beyond *ter* sites and their underlying mechanisms governing allowed and disallowed donors and insertion site pairings beyond preliminary observations will require the ability to perform systematic, high-throughput integration screens between many variant donor and recipient libraries in future studies. The ability of ACE to test the plasticity and requirement in the positional organization of information on the architectural genome provides an unique opportunity to comprehend, probe and reverse engineer the genomic underpinnings of life^5^.

In closing, we describe that ACE, and the iterations of ACE can integrate multi-megabase scale DNA across arbitrarily chosen donors and recipients to create artificial chimeric genomes. We further demonstrate that such chimeric genomes can be stable and can express the fused genomic pieces from multiple distinct parental species. With this novel ability, we echo the question first raised in Greek mythology: would it be possible to piece together a “chimeric” lifeform combining different biological traits across species, genus, or even order segregations? Robust and systematic generation of such “chimera” organisms would provide valuable insights on how the rationally designed synthetic chimeric genome directs the physiology and behavior of the underlying hybrid lifeform. Importantly, these techniques could lay the foundation of a new paradigm of artificial lifeforms to enable novel functions and future applications beyond the available designs of nature.

## Acknowledgement

We thank other members of the Kaihang Wang laboratory for advice and comments on the manuscript. We thank Prof. Takahito Mukai from Rikkyo University Japan for providing pBAD-traRP4min as a gift. We thank Prof. Shelley M Payne from the University of Texas at Austin for providing *Shigella flexneri* CFS100 as a gift. We thank Prof. Romain Koszul and his colleagues from Institute Pasteur France for providing the *Syn100* synthetic sequence. K.W. acknowledges funding support from the Shurl and Kay Curci Foundation, NIH (1DP2GM140937) and Caltech Center for Environmental Microbial Interactions (CEMI). R.J.Z. acknowledges support by the National Science Foundation Graduate Research Fellowship under Grant No. DGE-1745301

## Data and code availability

The sequencing data generated in this study will be submitted to the NCBI BioProject database and the accession number will be available before publication. A list of SRA accession numbers corresponding to each sequencing dataset is given in supplementary table X. Scripts used for sequencing data analysis are available upon request.

## Author contributions

C.S., R.J.Z. and K.W. developed the concept. C.S. and R.J.Z. performed experiments with help from G.W., M.W., and J. Huang, and analyzed data with input from K.W.. C.S., R.J.Z., J. Huang and M.L. developed Tn-Seq. C.S. developed sequencing data analysis scripts. J. Hur performed confocal microscopy. C.S., R.J.Z., G.W., M.W., M.L., J.Z. and R.S. built constructs. C.S., R.J.Z. and K.W. wrote the manuscript. All authors discussed the results and approved the manuscript.

## Declaration of interests

K.W. is a co-founder of Genyro Inc.. C.S., R.J.Z., Jianyi H., J.Z., K.W. are co-founders of Syntaxa Inc.. Neither company is involved in this study. A patent application on methods described in this paper has been filed by California Institute of Technology.

**Supplementary Figure 1.**
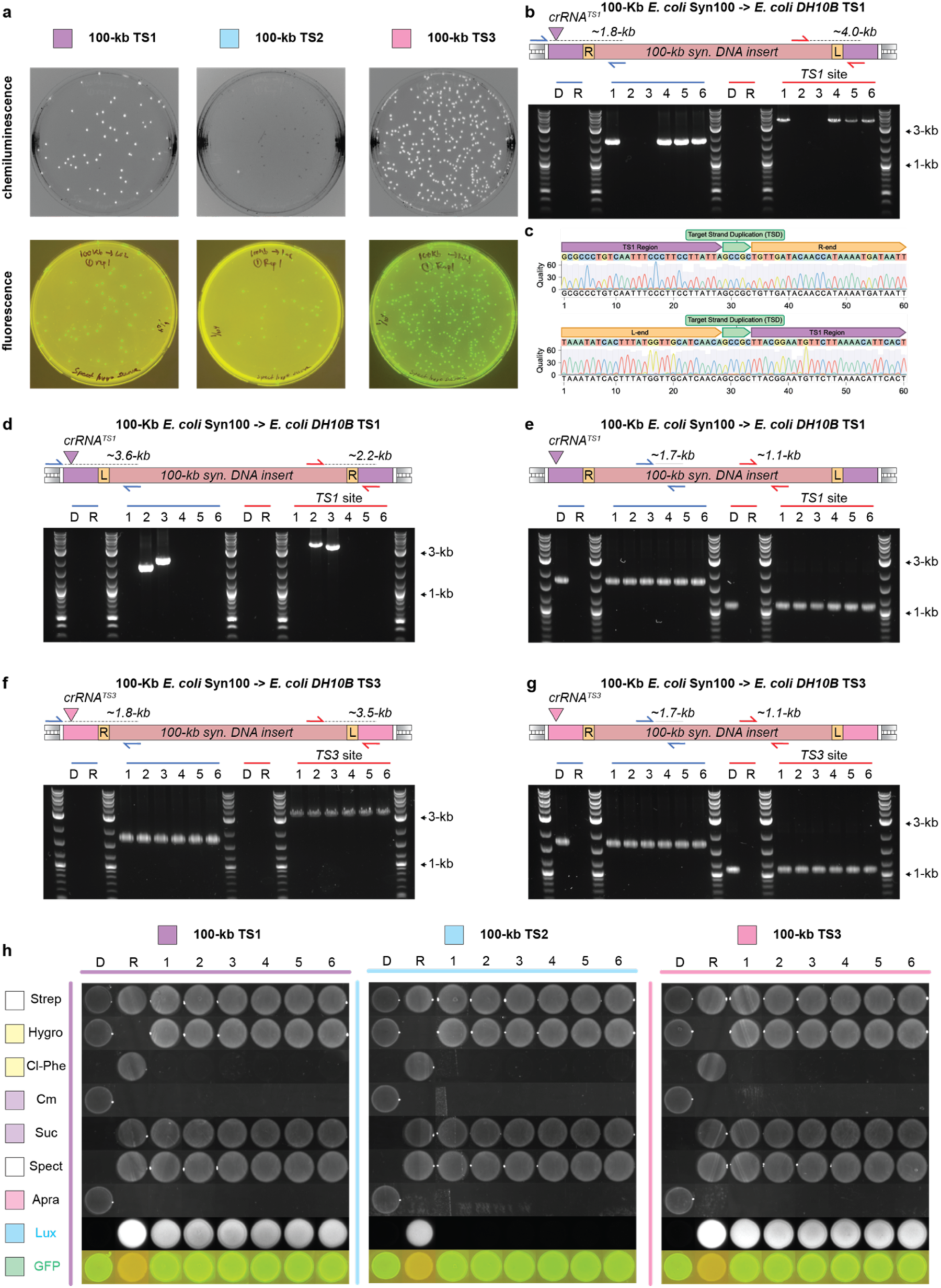
Detailed characterization of the fidelity and specificity of *Syn100* transfers with ACE. (**a**) Representative plates of 100-kb *Syn100* ACE transfers into TS1-3 in *E. coli* DH10B Δ*dif::lux*. The top panel shows colony luminescence while the bottom panel shows colony fluorescence under blue light. The 10^-4^ dilution plate was shown for each experiment. **(b**) Genotyping colony PCR gels for six integrant colonies of the 100-kb *Syn100* transfer into the DH10B Δ*dif::lux* TS1 site. In the schematic above the gel, the red rectangle represents the *Syn100* insert, the yellow boxes represent the LE and RE, and the purple rectangles represent the TS1 site. The integration site in TS1 targeted by crRNA*^TS1^* is shown with a purple triangle. The right and left junctions for a RL integration are checked with primers specific to TS1 and the *Syn100* segment, shown as blue and red arrows, respectively. D and R denote donor and recipient controls in each assay. Lengths of each PCR product are shown above each set of arrows in the schematic. **(c**) Sanger sequencing traces for the newly formed transposition junctions detected by genotyping colony PCR of clone 1 in b. for the 100-kb *Syn100* transfer into DH10B Δ*dif::lux*. **(d**) As in **b** but genotyping the same six integrant colonies for the right and left junctions for a LR integration, represented with the blue and red arrow pairs. **(e**) As in **b** but genotyping the same six integrant colonies for two internal stretches of the *Syn100* transfer with primer pairs denoted by the blue and red arrow pairs priming within the *Syn100* region. **(f**) As in **b** but genotyping colony PCR for six integrant colonies of the 100-kb *Syn100* transfer into DH10B Δ*dif::lux* TS3 site for the right and left junctions for a RL integration, represented with the blue and red arrow pairs. **(g**) As in **e** but genotyping the six integrant colonies of the 100-kb *Syn100* transfer into DH10B Δ*dif::lux* TS3 site for two internal stretches of the *Syn100* transfer with primer pairs denoted by the blue and red arrow pairs priming within the *Syn100* region. **(h**) Growth and fluorescence of *Syn100* donor (D), DH01B recipient (R) and the chimeric outcome strains (1-6) for each of the three integration positions (TS1-3) on different solid growth media conditions. Cells were spotted LB agar supplemented with one of the following in order from top to bottom (100 μg/ml streptomycin, 200 μg/ml hygromycin, 2.5 mM 4-chlorophenylalanine, 25 μg/ml chloramphenicol, 7.5% w/v sucrose, 50 μg/ml spectinomycin, 50 μg/ml apramycin). Chemiluminescence and fluorescence of the spotted colonies was assessed on the streptomycin condition.

**Supplementary Figure 2.**
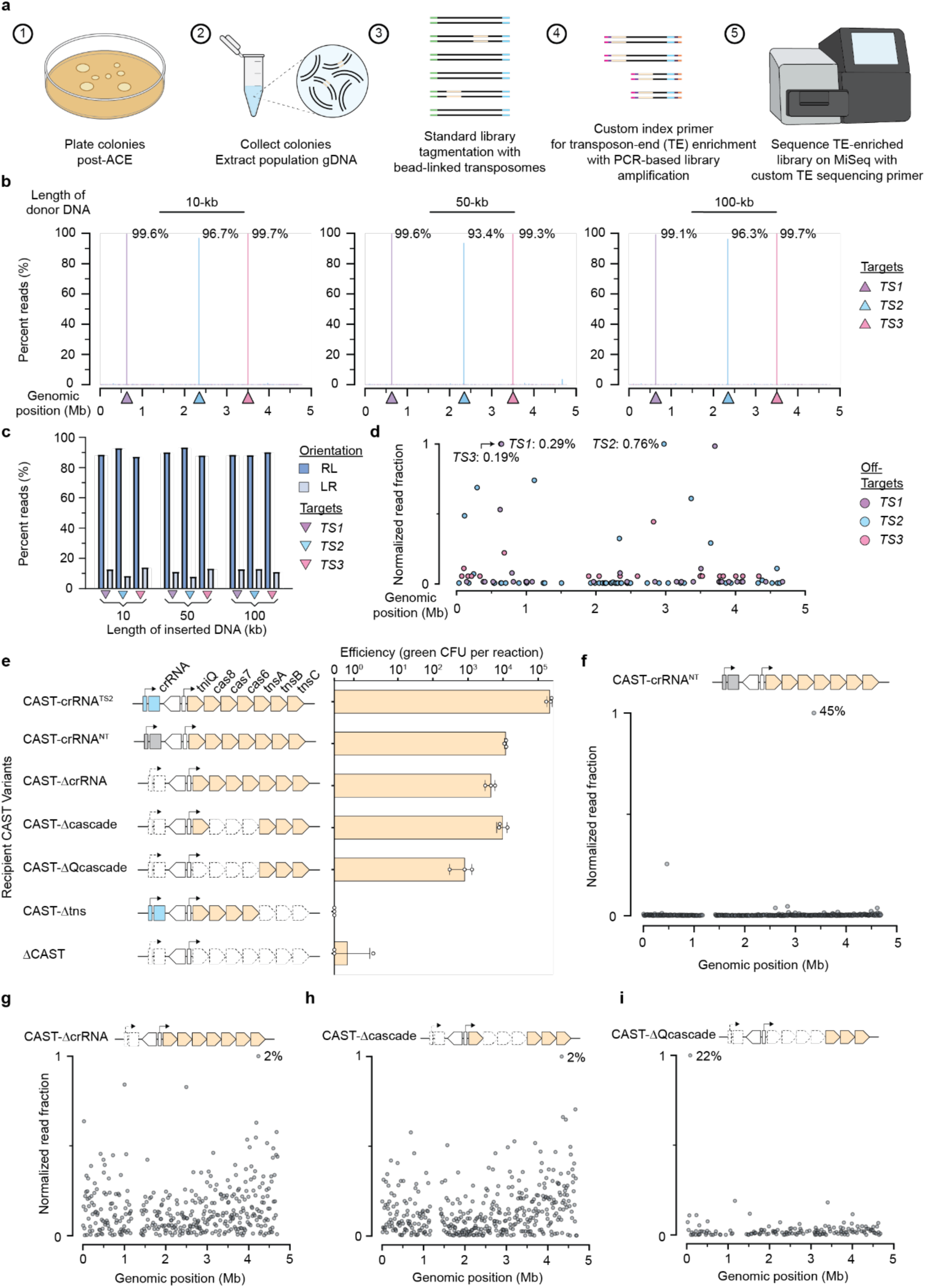
Description and demonstration of Tn-seq methodology and characterization of mutant CAST insertion behaviors with the *Syn100* donor. (**a**) Tn-seq workflow; (1) Bacterial colonies formed by ACE transfer on selective agar plates are (2) scraped and collected for genomic DNA extraction. A subpopulation of the extracted DNA contains fragments carrying transposon ends (TEs), denoted in beige, as a hallmark of a conjugative CAST event. (3) DNA fragments are randomly tagged with adapters and fragmented (tagmented) using bead-linked transposomes using routine library preparation protocols to append 5’ and 3’ sequences for standard sequencing primer annealing (blue and green). (4) Indices, P5 and P7 adapters are then added using a custom PCR-based library amplification step that allows for enrichment of sequences containing TEs from the total population of tagmented DNA. Briefly, custom primers containing the P5 adapter sequence (pink) as well as an index sequence (purple) were designed to anneal to the RE of the transposon (beige) were paired with a standard primer containing a P7 adapter (orange) and an index sequence (dark purple) which anneals to the standard sequencing primer binding site (blue) in a low cycle PCR. (5) This TE-enriched sequencing library is then sequenced on a MiSeq using a custom sequencing primer that anneals to the RE. **(b**) Genome-wide distribution of Tn-seq reads, measured in precent reads, from ACE transfer experiments of 10-kb, 50-kb and 100-kb *Syn100* cargoes into TS1 (purple bars), TS2 (blue bars) and TS3 (pink bars). Locations of each target site are given by triangles of the appropriate color. Genome coordinates are plotted with the origin of replication (*oriC*) as the zeroth coordinate. **(c**) Integration orientation for ACE transfer experiments of 10-kb, 50-kb and 100-kb *Syn100* cargoes into TS1 (purple), TS2 (blue) and TS3 (pink) given in percent Tn-seq reads. RL integration is given in blue and LR integration is given in light blue. **(d**) Genome-wide distribution of off-target Tn-seq reads for ACE transfer experiments of 100-kb *Syn100* cargo into TS1 (purple circles), TS2 (blue circles) and TS3 (pink circles). Frequency of off-target integration is given as a read fraction normalized to the most frequent off-target integration position. The most frequent off-target integration position for each of the TS1-3 experiments is additionally labeled with the percent reads in the Tn-seq mapping to said position. Genome coordinates are plotted with the origin of replication (*oriC*) as the zeroth coordinate. **(e**) Diagram illustrating the variants of the pAra-CAST plasmids used in 100-kb *Syn100* cargo transfers and corresponding efficiency of ACE transfer with each variant, measured in GFP+ c.f.u. Data is shown as the mean ± s.d. for *n* = 3 biologically independent samples. **(f**) Genome-wide distribution of Tn-seq reads for ACE transfer of 100-kb *Syn100* cargo with pAra-CAST-crRNA^NT^. Frequency of integration is given as a read fraction normalized to the most frequent integration position. The most frequent integration position is additionally labeled with the percent reads in the Tn-seq mapping to said position. Genome coordinates are plotted with the origin of replication (*oriC*) as the zeroth coordinate. **(g**) As in **f** but genome-wide distribution of Tn-seq reads for ACE transfer of 100-kb *Syn100* cargo with pAra-CAST-ΔcrRNA. **(h**) As in **f** but genome-wide distribution of Tn-seq reads for ACE transfer of 100-kb *Syn100* cargo with pAra-CAST-Δcascade. **(i**) As in **f** but genome-wide distribution of Tn-seq reads for ACE transfer of 100-kb *Syn100* cargo with pAra-CAST-ΔQcascade.

**Supplementary Figure 3.**
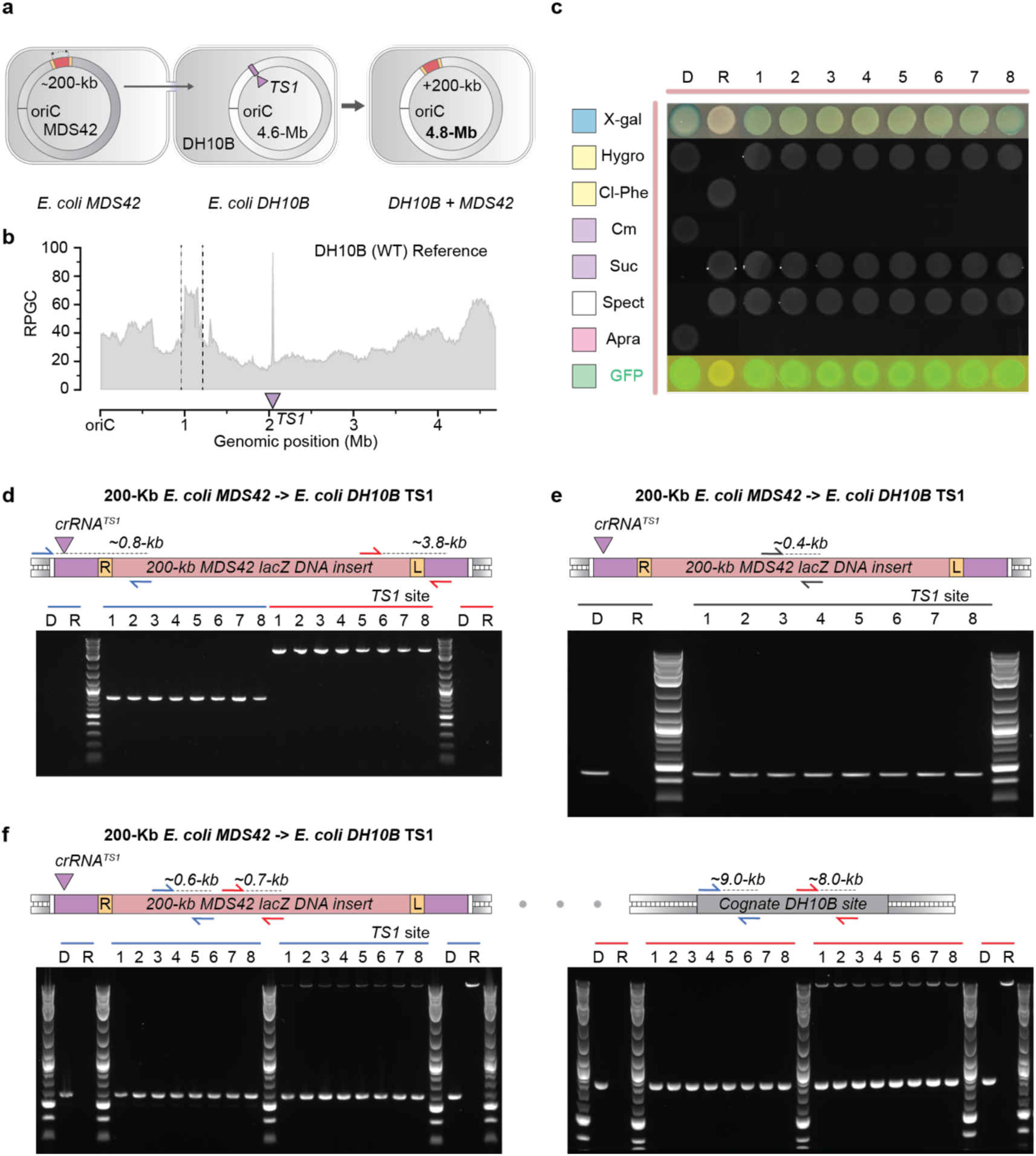
Description and validation of cross-strain, sub-megabase genomic DNA transfer in *E. coli*.. (**a**) Diagram of ∼200-kb *E. coli* MDS42 transfer to *E. coli* DH10B. The MDS42 genome is depicted by the dark gray circle and the DH10B genomes are depicted by the light gray circle. Sizes of transferred region, recipient genome and chimeric genome are given in the schematic. The transferred region in MDS42 is represented by the red portion of the circle flanked by TEs, shown as yellow rectangles. Transfer direction is shown by the arrow. The TS1 site of DH10B is given by the purple rectangle, which is targeted by the CAST by the TS1 crRNA, shown as a purple triangle. The outcome chimeric genome is represented as an integration of the red segment of the MDS42 genome into the DH10B genome at the TS1 site. **(b**) Genome-wide read coverages (in RPGC) for the *E. coli* DH10B + 200-kb MDS42 genomic chimeric strain. Region of addition is marked by dashed lines. Location of the TS1 target is given by the purple triangle. **(c**) Growth and fluorescence of MDS42 donor (D), DH01B recipient (R) and the chimeric outcome strains (1-8) on different solid growth media conditions. Cells were spotted LB agar supplemented with one of the following in order from top to bottom (20 μg/ml X-gal + 100 uM IPTG, 200 μg/ml hygromycin, 2.5 mM 4-chlorophenylalanine, 25 μg/ml chloramphenicol, 7.5% w/v sucrose, 50 μg/ml spectinomycin, 50 μg/ml apramycin, plain (-) LB agar). Fluorescence of the spotted colonies was assessed on the plain LB agar condition. (**d**) Genotyping colony PCR gels for eight integrant colonies of the 200-kb MDS42 transfer into the DH10B TS1 site. In the schematic above the gel, the red rectangle represents the 200-kb MDS42 insert, the yellow boxes represent the LE and RE, and the purple rectangles represent the TS1 site. The integration site in TS1 targeted by crRNA*^TS1^* is shown with a purple triangle. The right and left junctions for a RL integration are checked with primers specific to TS1 and the MDS42 segment, shown as blue and red arrows, respectively. D and R denote donor and recipient controls in each assay. Lengths of each PCR product are shown above each set of arrows in the schematic. **(e**) Genotyping colony PCR gels assaying for an internal segment of the MDS42 transfer with a primer pair denoted by gray arrows priming within the MDS42 region. Features of the integration schematic are as in **c**. **(f**) Genotyping colony PCR gels for eight integrant colonies of the 200-kb MDS42 transfer into the DH10B TS1 site using two primer pairs (denoted by the blue and red arrows, respectively) that hybridize within the integrated MDS42 segment in addition to a cognate region naturally in the DH10B genome, represented by a gray rectangle. In the first half of left gel, the blue primer pair is used in a PCR with a short ∼1-kb elongation time, while in the second half of the left gel, the blue primer pair used in a PCR with a long ∼10-kb elongation time. Likewise, in the first half of right gel, the red primer pair is used in a PCR with a short ∼1-kb elongation time, while in the second half of the right gel, the red primer pair used in a PCR with a long ∼10-kb elongation time. Features of the integration schematic are as in **c**.

**Supplementary Figure 4.**
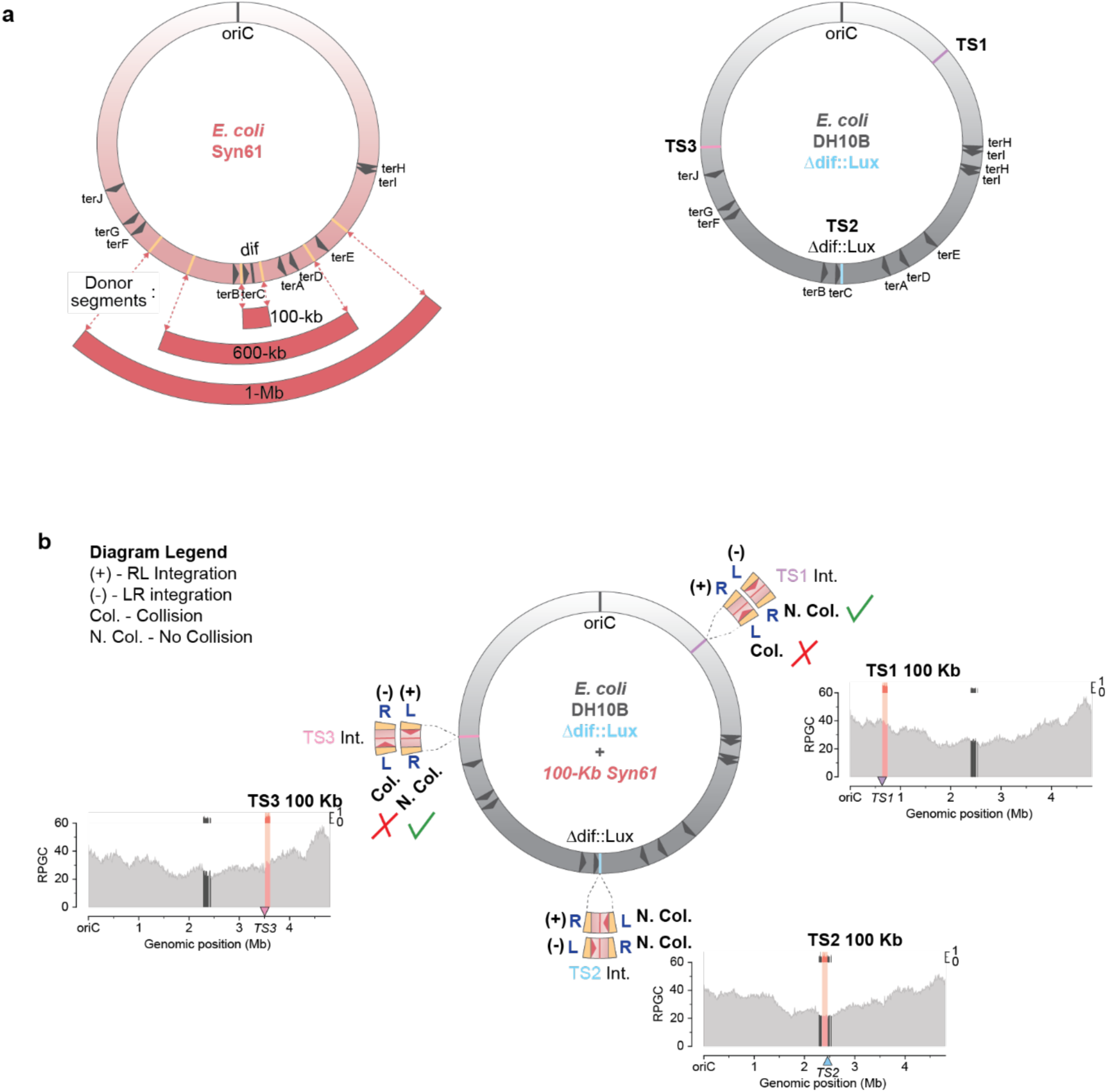
Orientation preference for 100 Kb Syn61 transfers into *E. coli* DH10B Δ*dif*::*lux*. (**a**) Diagram of the *E. coli* Syn61 (red) and *E. coli* DH10B Δ*dif*::*lux* (gray) genomes showing the positions of *ter* sites, *dif* site and origin of replication (*oriC*). Placements of the TEs are marked with yellow bars on the Syn61 genome corresponding to the 100-kb, 600-kb and 1-Mb Syn61 donor. Locations of the target sites 1-3 (TS1-3) in DH10B Δ*dif*::*lux* are represented with purple, blue and pink bars. **(b**) Integration diagram of DH10B Δ*dif*::*lux* genome with transfer of 100-kb Syn61 donor into either TS1, TS2 or TS3 via ACE. The transferred DNA is represented by the red ring segment flanked by yellow bars denoting the TEs labeled with either R or L to distinguish between RE and LE. Key sequences in the donor such as *ter* sites and the *dif* site are presented as red arrows and red bars, respectively. At each integration site (TS1-3), the two possible transposition orientations of the donor are depicted. (+) represents RL integration while (-) represents LR integration. At the TS1 and TS3 sites, one integration orientation and not the other would result in a chimeric genome that experiences collision (labeled Col. in the diagram) between replisome and the non-permissive side of a *tus*-bound *ter* site during chromosome synthesis, leading to termination of DNA replication (marked with a red X). The other orientation will lead to no collision (labeled N. Col.) as the replication fork will pass through the permissive end of the *tus-ter* complex (marked with a green checkmark). Adjacent to each integration is a corresponding genome-wide coverage map (in RPGC) for each *E. coli* DH10B + 100-kb Syn61 genomic chimera strain. Syn61 genomic segments are shaded in pink, and homologous *E. coli* DH10B genomic segments are shaded in dark gray. Non-redundant sections of the *E. coli* genome are in lighter grays. Locations of the TS is given by the triangle on the X-axis, triangles pointing down indicate the target is the sense strand, while triangles pointing up indicate the target is the antisense strand. Read fraction mapping to the Syn61 segment and the homologous DH10B segment is set above the coverage plot. Read fractions share the same color coding as the coverage plot.

**Supplementary Figure 5.**
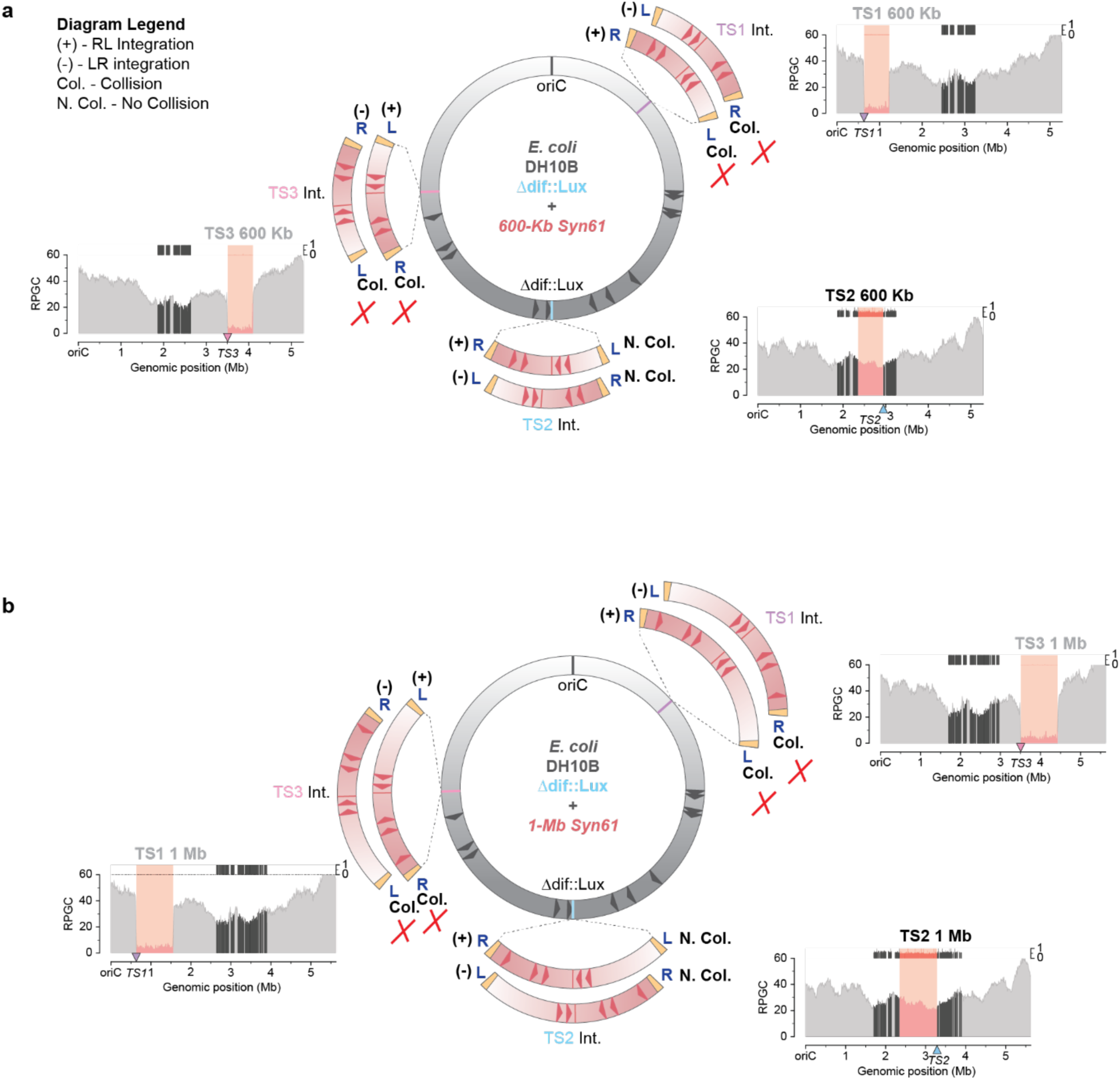
Architecture and stability of massively expanded *E. coli* synthetic genomes. (**a**) As in **Ext. Figure 4b** except integration diagrams of DH10B Δ*dif*::*lux* genome with transfer of 600-kb Syn61 donor into either TS1, TS2 or TS3 via ACE. At the TS1 and TS3 sites, both orientations of integration would result in replication fork collision with the non-permissive side of *tus*-bound *ter* site (Col., marked with a red X). Labels of TS1 and TS3 genome-wide coverage plots are grayed out to show instability and loss of the transferred segment following transposition. At the TS2 site, both integration orientations would lead to chimeric genomes that do not experience replication fork collision with the non-permissive side of a *ter*-*tus* complex. The corresponding genome coverage plot indicates the stability of the transferred segment. **(b**) As in **a** except integration diagrams of DH10B Δ*dif*::*lux* genome with transfer of 1-Mb Syn61 donor into either TS1, TS2 or TS3 via ACE.

**Supplementary Figure 6.**
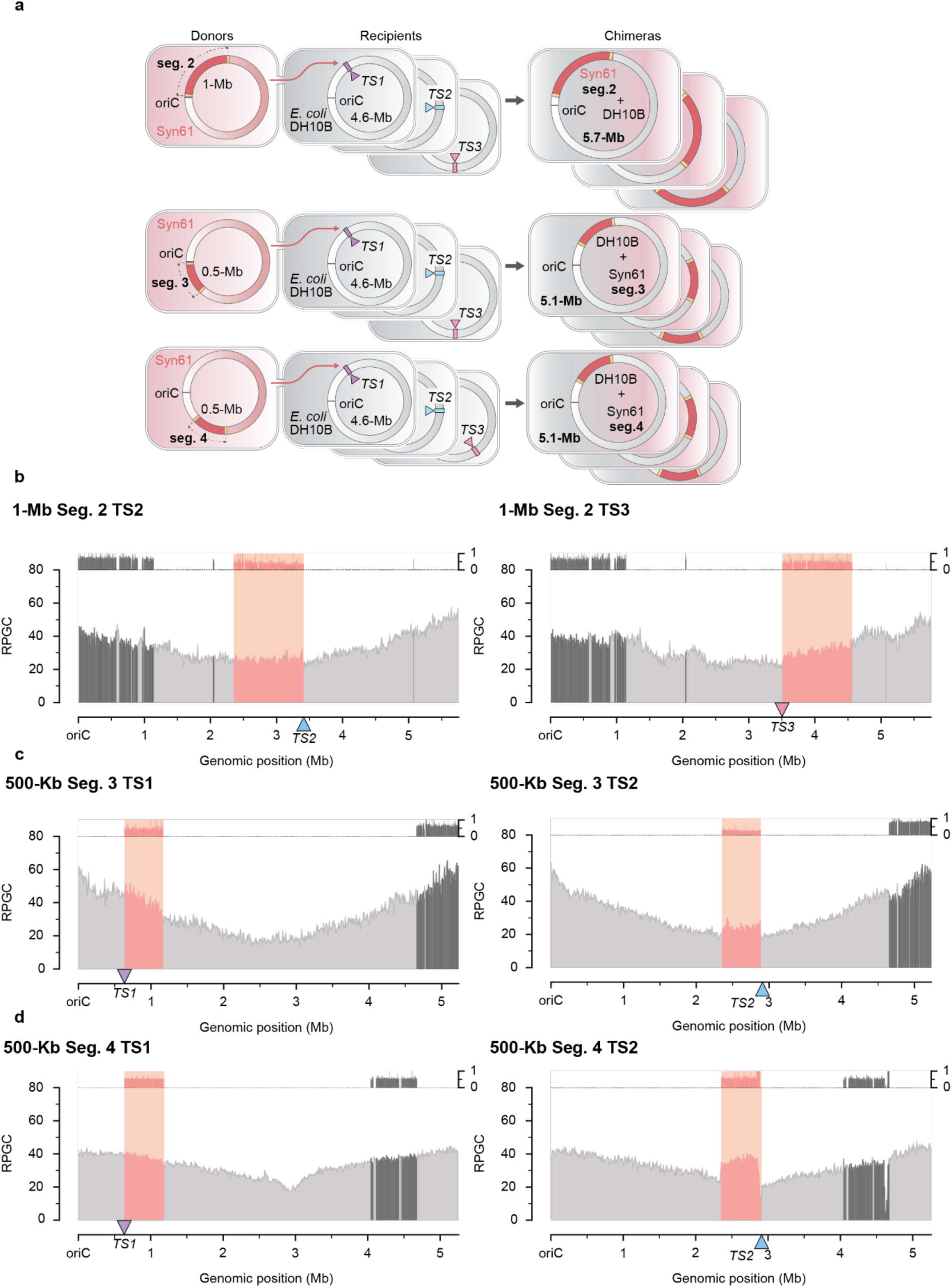
Stability and fidelity of Syn-segment transfers throughout the *E. coli* genome. (**a**) Diagrams representing the transfer of 1-Mb seg. 2, 500-kb seg. 3 and 500-Kb seg. 4 Syn61 donors for insertion at TS1, TS2 and TS3 in recipient *E. coli* DH10B *Δdif*::*lux* strains leading to nine possible chimeric genomes. The size of each outcome genome is labeled in bold in the chimeric strain. Note that integration orientation of the cargo is not explicitly depicted in the final product; each chimeric genome can adopt either a RL or LR configuration. **(b**) Genome-wide read coverages (in RPGC) for the *E. coli* DH10B + 1-Mb seg. 2 genomic chimeric strain integrated at TS2 and TS3. Syn61 genomic segments are shaded in pink, and homologous *E. coli* DH10B genomic segments are shaded in dark gray. Non-redundant sections of the *E. coli* genome are in lighter grays. Location of the TS2 and TS3 targets is given by the blue and pink triangles, respectively. Read fraction mapping to the Syn61 segment and the homologous DH10B segment is set above the coverage plot. Read fractions share the same color coding as the coverage plot. **(c**) As in **b** but genome-wide read coverage plot and read fraction mapping of 500-kb seg. 3 donor for insertion at TS1 (purple) and TS2 (blue) in recipient *E. coli* DH10B *Δdif*::*lux* strains. **(d**) As in **b** but genome-wide read coverage plot and read fraction mapping of 500-kb seg. 4 donor for insertion at TS1 (purple) and TS2 (blue) in recipient *E. coli* DH10B *Δdif*::*lux* strains.

**Supplementary Figure 7.**
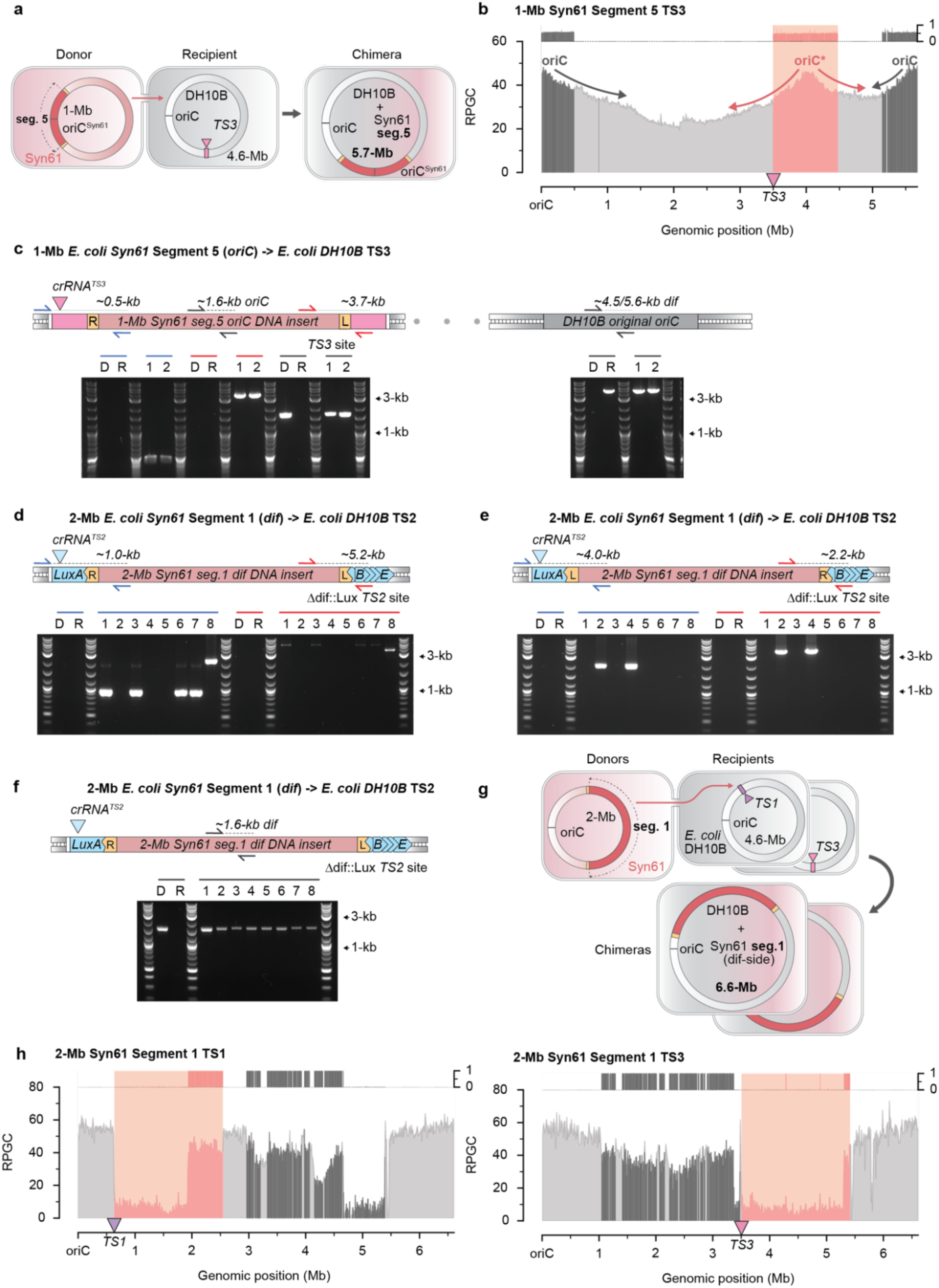
Installation of ectopic origin of replication in *E. coli* by means of Syn-segment 5 transfer and additional characterization of Syn-seg. 1 transfers. (**a**) Diagram of ∼1-Mb seg. 5 transfer to *E. coli* DH10B *Δdif*::*lux*. The Syn61 genome is depicted by the pink circle and the *E. coli* DH10B genome are depicted by the gray circle. Sizes (in Mb) of transferred region, recipient genome and chimeric genome are given in the schematic. The transferred region is centered around the *oriC* of Syn61 and is represented by the red portion of the circle flanked by TEs, shown as yellow rectangles. Transfer direction is shown by the red arrow. The TS3 site of *E. coli* DH10B *Δdif*::*lux* is given by the pink rectangle, which is targeted by the CAST by crRNA*^TS2^*, shown as a pink triangle. The outcome chimeric genome (5.7 Mb) is represented as an integration of the red segment of the Syn61 genome into the gray DH10B genome at the TS3 site. **(b**) Genome-wide read coverages (in RPGC) for the *E. coli* DH10B + 1-Mb Syn-Segment 5 genomic chimeric strain integrated at TS3. Syn61 genomic segments are shaded in pink, and homologous *E. coli* DH10B genomic segments are shaded in dark gray. Non-redundant sections of the *E. coli* genome are in lighter grays. Location of the TS3 target is given by the blue triangle. The genomic chimera produced by this transfer has an ectopic *oriC* inserted at the TS3 region, which forms an additional peak in read coverage alongside the wild type *oriC*. Dark gray and pink arrows illustrate the two sets of replication forks emanating from the WT and ectopic *oriC*s, respectively. Read fraction mapping to the Syn61 segment and the homologous DH10B segment is set above the coverage plot. Read fractions share the same color coding as the coverage plot. **(c**) Genotyping colony PCR gels for two integrant colonies of the 1-Mb seg. 5 transfer into *E. coli* DH10B TS3. In the schematic above the gel, the red rectangle represents the 1-Mb seg. 5 insert, the yellow boxes represent the LE and RE, and the pink rectangles represent the TS3 site. The integration site is targeted by crRNA*^TS3^* shown with a pink triangle. The right and left junctions for a RL integration are checked with primers specific to TS3 and 1-Mb seg. 5, shown as blue and red arrows, respectively. The Syn61 *oriC* and the DH10B *oriC* are also assayed with specific primers for each locus as with the black arrows. D and R denote donor and recipient controls in each assay. Lengths of each PCR product are shown above each set of arrows in the schematic. **(d**) Genotyping colony PCR gels for eight integrant colonies of the 2-Mb seg. 1 transfer into the *E. coli* DH10B *Δdif*::*lux* TS2 site. In the schematic above each gel, the red rectangle represents the 2-Mb seg. 1 insert, the yellow boxes represent the LE and RE, and the blue squares rectangles and triangles represents the *lux* operon. The integration site in *luxA* targeted by crRNA*^TS2^* is shown with a blue triangle. The right and left junctions for a RL integration are checked with primers specific to TS2 and seg. 1, shown as blue and red arrows, respectively. D and R denote donor and recipient controls in each assay. Lengths of each PCR product are shown above each set of arrows in the schematic. **(e**) As in **d** but genotyping the same eight integrant colonies for the right and left junctions for a LR integration, represented with the blue and red arrow pairs. **(f**) As in **d** but genotyping the same eight integrant colonies for Syn61 *dif* site with a primer pair denoted by the black arrow pair priming within the Syn61 *dif* region. **(g**) Diagrams representing the transfer of 2-Mb seg. 1 Syn61 donors for insertion at TS1 and TS3 in recipient *E. coli* DH10B *Δdif*::*lux* strains. The size of the outcome genome is labeled in bold in the chimeric strain. Note that integration orientation of the cargo is not explicitly depicted in the final product; each chimeric genome can adopt either a RL or LR configuration. **(h**) Genome-wide read coverages (in RPGC) for the *E. coli* DH10B + 2-Mb seg. 1 genomic chimeric strain integrated at TS1 and TS3. Syn61 genomic segments are shaded in pink, and homologous *E. coli* DH10B genomic segments are shaded in dark gray. Non-redundant sections of the *E. coli* genome are in lighter grays. Location of the TS1 and TS3 targets is given by the purple and pink triangles, respectively. Read fraction mapping to the Syn61 segment and the homologous DH10B segment is set above the coverage plot. Read fractions share the same color coding as the coverage plot.

**Supplementary Figure 8.**
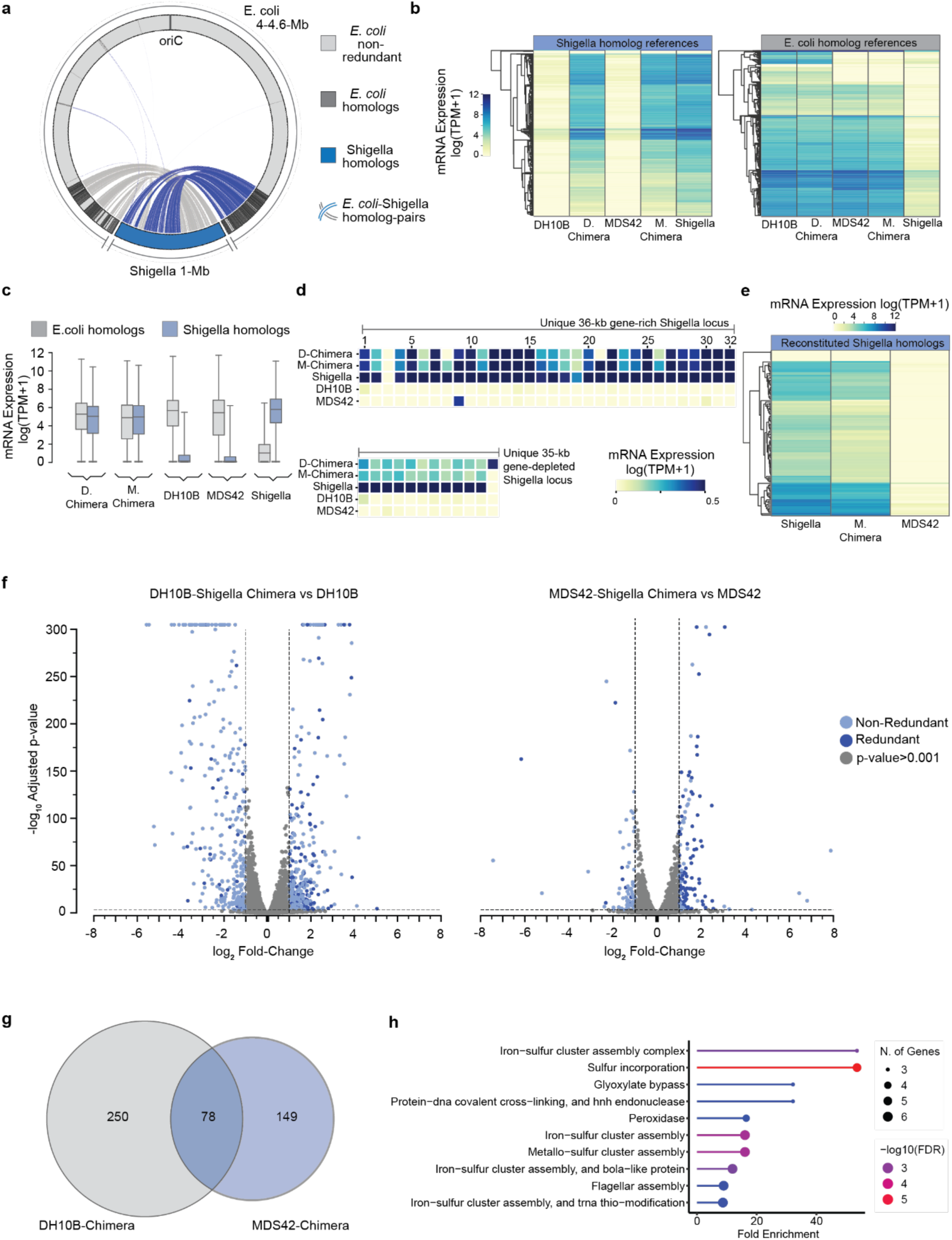
Transcriptomic characterization of *E. coli Shigella* genomic chimeras. (**a**) Genome-wide mapping of *S. flexneri* gene variants given in blue to their corresponding *E. coli* wild-type homologs given in dark gray in the *E. coli* + 1-Mb *S. flexneri* chimeric genome with a R-L integration orientation. Pairs of DH10B and *Shigella* homologs are connected by either blue or light gray lines. **(b**) Hierarchical clustering of transcript counts, measured in log of transcripts per million + 1 (log(TPM + 1)), of *S. flexneri* gene homologs and *E. coli* gene homologs in the DH10B, MDS42 DH10B*-Shigella*, MDS42*-Shigella* and *Shigella* strains. Three individual replicates are shown for each strain. Genes are clustered based on expression level within each genomic background. **(c**) Expression distribution of bulk *S. flexneri* transcripts (blue) compared to bulk homologous *E. coli* transcripts (gray) in DH10B, MDS42, *S. flexneri,* and Chimeric strains measured in log(TPM+1). Distributions are based on the data from *n = 3* biologically independent samples. Edges of boxes close to zero represent outliers that are the result of bioinformatic and sequencing artifacts with no biological significance. **(d**) Positional heat map of expression in log(TPM+1) of two unique gene-rich *S. flexneri* loci in DH10B, MDS42, *S. flexneri*, and their respective *E. coli* + 1-Mb *S. flexneri* chimeras. **(e**) Hierarchical clustering of transcript counts, measured in log(TPM+1) in the *S. flexneri*, Chimeric and MDS42 strains, of *S. flexneri* gene homologs originally deleted in the MDS42 strain background that are reconstituted via ACE chimerization with *S. flexneri*. Three individual replicates are shown for each strain. Genes are clustered based on expression level within each genomic background. **(f**) Differential expression of redundant genes (dark blue) and non-redundant genes (light blue) between the chimeric strain and the *E. coli* background strain for both *E. coli* DH10B + 1 Mb *S. flexneri and E. coli* MDS42 + 1 Mb *S. flexneri*. The log_2_ of the ratio of the expression level of the redundant genes in the chimeric strain to the expression level of their homolog in the *E. coli* strain (dark blue) is plotted alongside log_2_ of the ratio of the expression level of non-redundant genes in the chimeric strain to the expression level of its homolog in the *E. coli* strain (light blue). Points are also plotted based on their –log_10_ adjusted p-value. P-value cutoff was set at 0.001 and the log_2_ fold-change cutoff was set to ±1 so that points below the cutoff are in gray. **(g**) Venn diagram of the redundant genes overexpressed in the *E. coli* DH10B + 1-Mb *S. flexneri* chimera and the *E. coli* MDS42 + 1-Mb *S. flexneri* chimera and highlighting the set of redundant genes present and overexpressed in both chimeric strains. **(h**) Analysis of the set of redundant genes overexpressed in both chimeric strains, grouped by GO (gene ontology) terms. Functional groupings are arranged by fold-enrichment of overexpression as well as the –log_10_ false discovery rate (FDR).

**Supplementary Figure 9.**
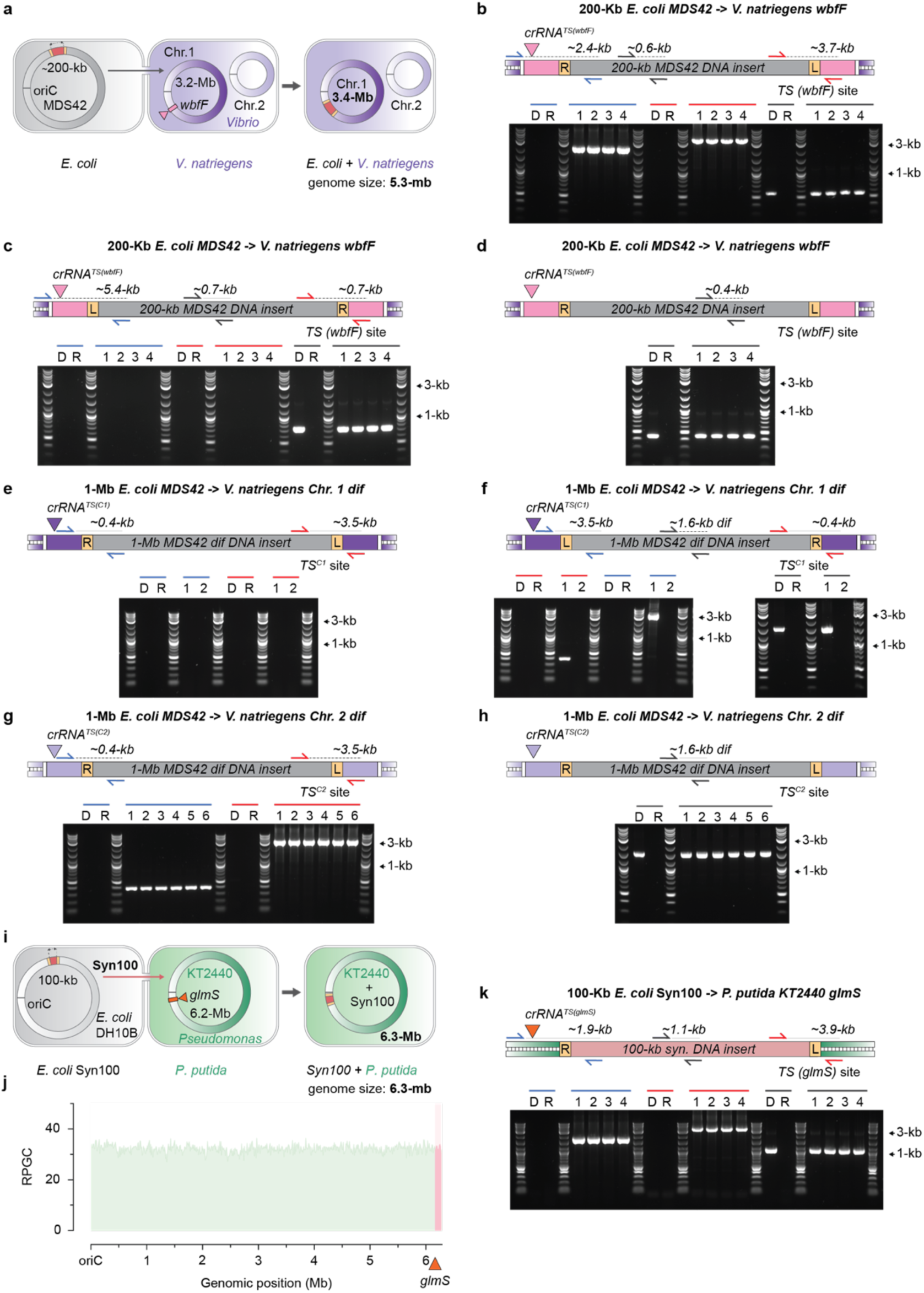
Genomic transfers from *E. coli* to *V. natriegens* and *P. putida*. (**a**) Diagram of 200-kb *E. coli* MDS42 transfer to *V. natriegens*. The MDS42 genome is depicted by the gray circle and the *V. natriegens* genomes are depicted by the purple circles, with chromosome 1 in dark purple and chromosome 2 in light purple. Sizes of transferred region, recipient genome and chimeric genome are given in the schematic. The transferred region in MDS42 is represented by the red portion of the circle flanked by TEs, shown as yellow rectangles. Transfer direction is shown by the arrow. The *wbfF* gene of *V. natriegens* chromosome 1 is given by the pink rectangle, which is targeted by the CAST by the TS*^wbfF^* crRNA, shown as a pink triangle. The outcome chimeric genome is represented as an integration of the red segment of the MDS42 genome into the *V. natriegens* genome at *wbfF*. **(b**) Genotyping colony PCR gels for four integrant colonies of the 200-kb MDS42 transfer into the *V. natriegens wbfF*. In the schematic above the gel, the gray rectangle represents the 200-kb MDS42 insert, the yellow boxes represent the LE and RE, and the pink rectangles represent the *wbfF* site. The integration site in *wbfF* targeted by crRNA*^wbfF^* is shown with a pink triangle. The right and left junctions for a RL integration are checked with primers specific to *wbfF* and the MDS42 segment, shown as blue and red arrows, respectively. An internal region of the MDS42 integration was also checked, shown with the gray arrows. D and R denote donor and recipient controls in each assay. Lengths of each PCR product are shown above each set of arrows in the schematic. **(c**) As in **b** but genotyping colony PCR for the four integrant colonies for the right and left junctions for a LR integration, represented with the blue and red arrow pairs, as well as for another internal region along the MDS42 segment given with the gray arrows. **(d**) As in **b** but with an additional genotyping PCR check of a region within the integrated MDS42 segment in the *wbfF* site of *V. natriegens*. **(e**) Genotyping colony PCR gels for two integrant colonies of the 1-Mb MDS42 transfer into the *V. natriegens* chromosome 1 *dif*. In the schematic above the gel, the gray rectangle represents the 1-Mb MDS42 insert, the yellow boxes represent the LE and RE, and the dark purple rectangles represent the chr. 1 *dif* site. The integration site is targeted by crRNA*^TS(C1)^* shown with a dark purple triangle. The right and left junctions for a RL integration are checked with primers specific to chr. 1 *dif* and the MDS42 segment, shown as blue and red arrows, respectively. D and R denote donor and recipient controls in each assay. Lengths of each PCR product are shown above each set of arrows in the schematic. **(f**) As in **c** but genotyping colony PCR for the two integrant colonies for the right and left junctions for a LR integration, represented with the blue and red arrow pairs, as well as for an internal region along the MDS42 segment represented with the gray arrows. **(g**) Genotyping colony PCR gels for two integrant colonies of the 1-Mb MDS42 transfer into the *V. natriegens* chromosome 2 *dif*. In the schematic above the gel, the gray rectangle represents the 1-Mb MDS42 insert, the yellow boxes represent the LE and RE, and the light purple rectangles represent the chr. 2 *dif* site. The integration site is targeted by crRNA*^TS(C2)^*shown with a light purple triangle. The right and left junctions for a RL integration are checked with primers specific to chr. 2 *dif* and the MDS42 segment, shown as blue and red arrows, respectively. D and R denote donor and recipient controls in each assay. Lengths of each PCR product are shown above each set of arrows in the schematic. **(h**) Genotyping colony PCR gels assaying for an internal segment of the MDS42 transfer with a primer pair denoted by gray arrows priming within the MDS42 region. Features of the integration schematic are as in **e**. **(i**) Diagram of 100-kb *Syn100* transfer to *Pseudomonas putida*. The *E. coli Syn100* genome is depicted by the gray circle and the *P. putida* genome is depicted by the green circle. Sizes of transferred region, recipient genome and chimeric genome are given in the schematic. The transferred region *Syn100* is represented by the red portion of the circle flanked by TEs, shown as yellow rectangles. Transfer direction is shown by the arrow. The *glmS* gene of *P. putida* is given by the orange rectangle, which is targeted by the CAST by the TS*^glmS^*crRNA, shown as an orange triangle. The outcome chimeric genome is represented as an integration of the red *Syn100* segment into the *P. putida* genome at *glmS*. **(j**) Genome-wide read coverages (in RPGC) for the *P. putida* KT2440 + 100-kb *Syn100* genomic chimeric strain. The *Syn100* segment is shaded in red, and *P. putida* genome is shaded in light green. Location of the TS *glmS* target is given by the orange triangle. **(k**) Genotyping colony PCR gels for four integrant colonies of the 100-kb *Syn100* transfer into the *P. putida glmS*. In the schematic above the gel, the red rectangle represents the 100-kb *Syn100* insert, the yellow boxes represent the LE and RE, and the green duplex DNA represents the *glmS* site. The integration site in *glmS* targeted by crRNA*^glmS^* is shown with an orange triangle. The right and left junctions for a RL integration are checked with primers specific to *glmS* and the *Syn100* segment, shown as blue and red arrows, respectively. An internal region of the *Syn100* integration was also checked, shown with the gray arrows. D and R denote donor and recipient controls in each assay. Lengths of each PCR product are shown above each set of arrows in the schematic.

**Supplementary Figure 10.**
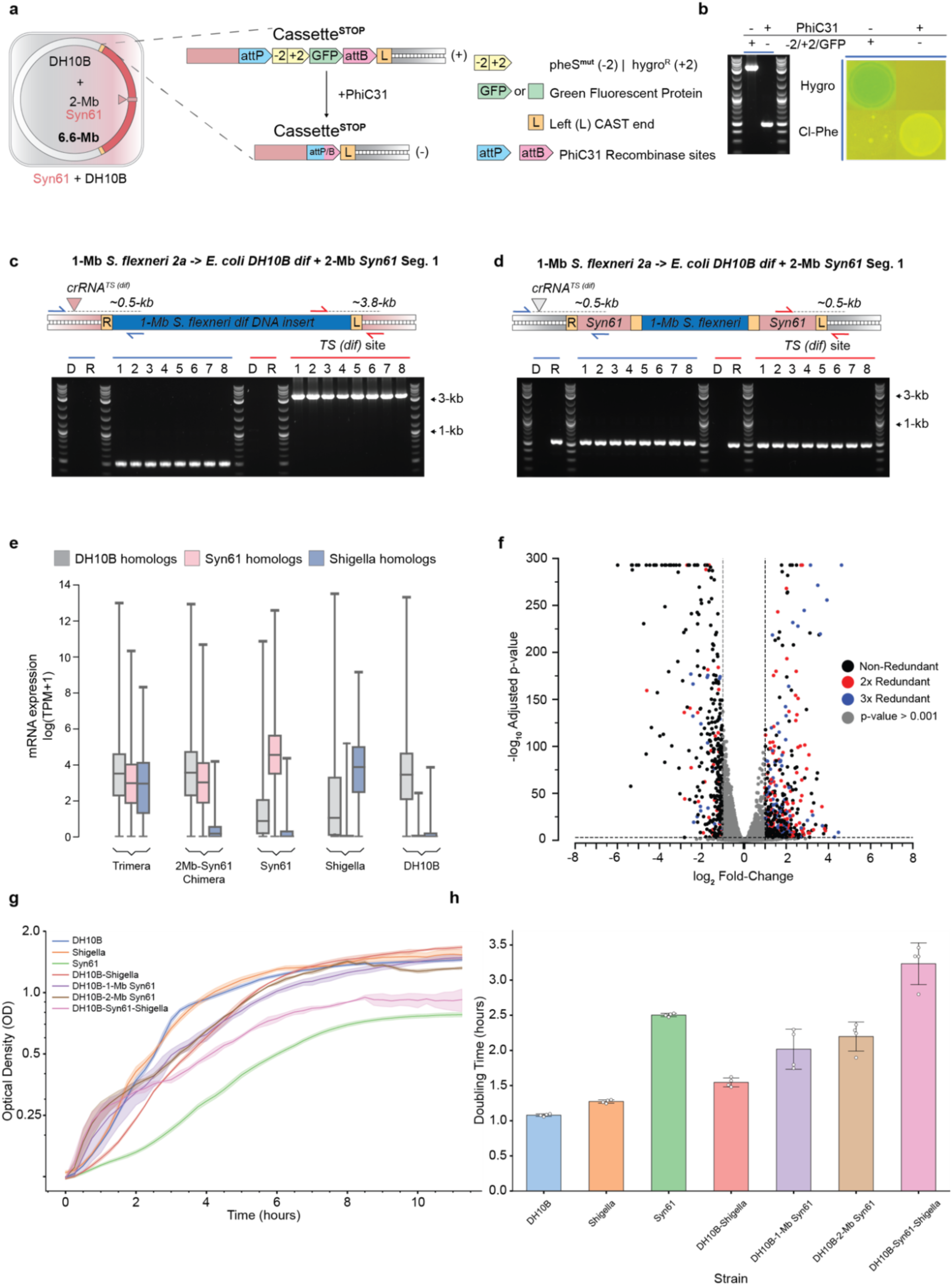
Construction, validation, and characterization of an organism with a trimeric genome. (**a**) Using PhiC31 recombinase to excise +2/2 selection and GFP markers from Cassette*^END^* of a *dif* integrated *E. coli* DH10B + 2-Mb Syn-seg. 1 genome chimera strain. The expression of *PhiC31* in this strain triggers a crossing-over event between the *attP* and *attB* sequences flanking the selection and screening markers in Cassette*^END^*, leading to marker loss, and enabling subsequent ACE events in this chimeric strain as shown in **Figure 5e**. **(b**) Assaying for Cassette*^END^* marker excision via genotyping colony PCR of *dif* integrated *E. coli* DH10B + 2-Mb seg. 1 cells before (+) and after (-) expression of *PhiC31* recombinase. Chimeric cells before (+) and after (-) *PhiC31* induction were also stamped onto selective LB agar containing either 200 μg/ml hygromycin (Hygro) or 2.5 mM 4-chlorophenylalanine (Cl-Phe). **(c**) Genotyping colony PCR gels for eight integrant colonies of the 1-Mb *S. flexneri* transfer into the *dif* integrated *E. coli* DH10B + 2-Mb seg. 1. In the schematic above the gel, the blue rectangle represents the 1 Mb *Shigella* insert, the yellow boxes represent the LE and RE, and the red duplex DNA represents 2 Mb seg. 1 *dif* site. The integration site in *dif* targeted by crRNA*^TS(dif)^* is shown with a red triangle. The right and left junctions for a RL integration are checked with primers specific to *dif* and the *Shigella* segment, shown as blue and red arrows, respectively. D and R denote donor and recipient controls in each assay. Lengths of each PCR product are shown above each set of arrows in the schematic. **(d**) Genotyping colony PCR gels for eight integrant colonies of the 1-Mb *S. flexneri* transfer into the *dif* integrated *E. coli* DH10B + 2-Mb Syn-seg. 1. In the schematic above the gel, the blue rectangle represents the 1 Mb *Shigella* insert, the yellow boxes represent the LE and RE, the red rectangles represent 2-Mb seg. 1 genomic stretches and the gray duplex DNA represents the DH10B *dif* site. The integration site in *dif* targeted by crRNA*^TS(dif)^* is shown with a gray triangle. The right and left junctions for a RL integration of 2-Mb seg. 1 at the DH10B *dif* are checked with primers specific to *dif* and the Syn61 segment, shown as blue and red arrows, respectively. D and R denote donor and recipient controls in each assay. Lengths of each PCR product are shown above each set of arrows in the schematic. **(e**) Expression distribution of bulk Syn61 transcripts (pink), *S. flexneri* transcripts (blue) compared to bulk homologous DH10B transcripts (gray) in DH10B, *S. flexneri*, Syn61, Chimeric and Trimeric strains measured in log(TPM+1). Distributions are based on the data from *n = 3* biologically independent samples. Edges of boxes close to zero represent outliers that are the result of bioinformatic and sequencing artifacts with no biological significance. **(f**) Differential expression of 3-fold (blue), 2-fold redundant genes (red) and non-redundant genes (black) between the trimeric strain and the DH10B background strain. The log_2_ of the ratio of the expression level of the 3-fold and 2-fold redundant genes in the trimeric strain to the expression level of their homolog in the DH10B strain (blue and red) is plotted alongside log_2_ of the ratio of the expression level of non-redundant genes in the trimeric strain to the expression level of its homolog in the DH10B strain (black). Points are also plotted based on their –log_10_ adjusted p-value. P-value cutoff was set at 0.001 and the log_2_ fold-change cutoff was set to ±1 so that points below the cutoff are in gray. **(g**) Semi-log plot of growth curves (optical density as a function of hours elapsed) for chimeric and trimeric genome bacteria as well as their wild-type precursors strains. *E. coli* DH10B is given by the blue curve, *S. flexneri* is given by the orange curve, *E. coli* Syn61 is given by the green curve, DH10B + 1-Mb *Shigella* is given by the red curve, DH10B + 1-Mb Syn61 is given by the purple curve, DH10B + 2-Mb Syn61 is given by the brown curve and DH10B + 2-Mb Syn61 + 1-Mb *Shigella* is given by the pink curve. Reported curves are the average of four independently grown biological replicates of each strain, with the solid curve representing the mean OD at each time point and the colored halo surrounding each curve representing +/-s.d. Note that optical density is plotted on a logarithmic scale, hence the semi-log plot. **(h**) Doubling times for chimeric and trimeric genome bacteria as well as their wild-type precursor strains. Reported doubling time, shown as mean +/− s.d., are the average of four independently grown biological replicates of each strain. The doubling time reported for each individual experiment is represented by dots.

## Methods

### General methods

DNA amplification was performed using PrimeSTAR^TM^ GXL DNA Polymerase (TaKaRa Bio USA) or Q5^®^ High-Fidelity 2x Master Mix (New England BioLabs (NEB)) unless otherwise stated. DNA oligonucleotides were obtained from Integrated DNA Technologies (IDT) and Millipore Sigma. Gene fragments (gBlocks) were obtained from IDT or Twist Biosciences. Nucleic acid purification and cleanup was done with QIAquick^®^ PCR Purification Kit (QIAGEN^™^ cat # 28106). DNA extraction from agarose gels was conducted with GeneJET Gel Extraction Kit (Thermo Scientific™ cat # K0691). All plasmids were assembled using Gibson assembly using NEBuilder^®^ 2x HiFi Assembly mix (NEB). Standard plasmid DNA cloning was performed using electrocompetent *E. coli* DH10B, which was made electrocompetent as previously described^1^. Routine of plasmid DNA extraction from *E. coli* DH10B was done with QIAprep^®^ Spin Miniprep Kit (QIAGEN^™^ cat # 27106). All antibiotic and counter-selection concentrations when used are as follows: 100 μg/ml streptomycin, 200 μg/ml hygromycin, 2.5 mM 4-chlorophenylalanine, 25 μg/ml chloramphenicol, 7.5% w/v sucrose, 50 μg/ml spectinomycin, 50 μg/ml apramycin.

### Bacteria culturing conditions

All *E. coli* and *S. flexneri* bacterial strains were grown at 37°C in selective Luria-Bertani (LB) liquid media with shaking unless otherwise stated. Individual colonies were grown overnight on selective LB-agar plates at 37 °C prior to liquid growth. All *P. putida* bacterial strains were grown at 30°C in LB liquid media with shaking or on solid LB-agar plates. For overnight culturing, *V. natriegens* bacterial strains were grown at 30°C in LBv2 media for liquid growth or on LB-agar plates for solid growth. For daytime culturing for electrocompetency, *V. natriegens* was grown at 37°C in LBv2 media. *V. natriegens* electrocompetency was performed as previously described^2^.

Detailed documentation of all bacteria strains and their derivatives used in this study are listed in Supplemental Table 1. Briefly, *E. coli* DH10B *rpsL K43R*, *E. coli* DH10B *rpsL K43R* Δ*dif*::*lux*, *E. coli* MDS42 *rpsL K43R ΔrecA*, *Vibrio natriegens* ATCC 14048 DSM759 and *Pseudomonas putida* KT2440 were used as base strains for the construction of ACE recipient strains. Donor strain construction was carried out in *E. coli* DH10B *rpsL K43R* Syn100, *E. coli* MDS42 *rpsL K43R ΔrecA*, *E. coli* MDS42 *rpsL K43R ΔrecA Δtus*, *E. coli* Syn61 *rpsL K43R ΔpheS*-hygroR* and *Shigella flexneri* CFS100 *rpsL K43R* Δ*recA* Δ*tus* Δ*oriT* Δ*higB* Δ*hok* backgrounds.

### Plasmid construction

All plasmids and oligonucleotides used in their construction are listed in Supplemental Table 2 and 3. All plasmid constructs described in this section were constructed via Gibson assembly. In short, *pAra-CAST* was derived from pSL1777 (Addgene #160731). *pAra-CAST* variants under the RSF backbone were constructed for experiments carried out in *V. natriegens* and *P. putida*. The conjugation component used in this work (*pAra-Trans*) was derived from pBAD-traRP4min and the commercial pBelo ori2 BAC. Plasmid templates for Cassette^START^ (*pSC101-spectR-cassette1-sC-oriT-RE*) and Cassette^END^ (*pSC101-spectR-cassette2-pHGFP-LE-oriT*) were cloned under a pSC101 backbone to facilitate propagation for downstream amplification and λ-red mediated recombination.

### Strain construction

Classical λ-red mediated homologous recombination was also used to integrate Cassette^START^ and Cassette^END^ into the genome to generate donor strains^3,4^. Briefly, pCassette^START^ was amplified with PCR using primer pairs bearing homology to the start and end of the transferred region and transformed into pre-induced electrocompetent cells of the relevant bacterial strain^1^ (see Supplemental Table 3 for primer pairs used). Cells were plated on selective media and genotyped by PCR for cassette integration with the relevant primers in Supplemental Table 3. Colonies with the correct integration were sequenced. Cassette^END^ was integrated into colonies with sequence verified Cassette^START^ using the same procedure. Strains with both Cassettes integrated into their genome were made electrocompetent for transformation of *pAra-Trans* by electroporation as previously described to make finalized donor strains. Likewise, to generate recipient strains, base bacterial strains were made electrocompetent and transformed with *pAra-CAST* by electroporation to finalize recipient strains.

### Conjugative-transposition experiments

All ACE events were performed at 30°C based on past reports that CAST-directed transposition is more efficient at 30°C than at 37°C.

For ACE transfer experiments involving *E. coli DH10B* Syn100, *S. flexneri*, *V. natriegens* or *P. putida* donor strains, donor and recipient cultures were inoculated into 10 mL of LB media with appropriate antibiotics and 2% (w/v) glucose for overnight growth (12-16 hrs). Overnight cultures were then pelleted by centrifugation and diluted into 100 mL of LB with the appropriate antibiotics at a starting OD_600_ of 0.05. Cultures were grown at 37°C to OD_600_ of 0.10-0.20 before inducing with 0.5% (w/v) arabinose until cultures reach an OD_600_ of 0.5. Cultures were then pelleted and resuspended in 400 uL of LB. Resuspended donors and recipients were then mixed in a 4:1 donor to recipient ratio (320 uL + 80 uL) for a total of 400 uL and spotted in 13 uL aliquots onto a LB-agar plate with no antibiotics. Spots were dried under a flame before incubating at 30°C for 4-8 hours to allow for conjugation coupled transposition activity. Spots were then washed from the plate using 5 mL of LB and added to a culture 95 mL of LB with the appropriate antibiotics and 0.5% (w/v) arabinose. Cultures were recovered at 37°C for 3-6 hours before pelleting by centrifugation and then plating by serial dilution on selective LB-agar plates with appropriate antibiotics. Plates were incubated at 30°C for 12-16 hours before shifting up to 37°C for another 12 hours to accelerate colony growth, except for transfers into *P. putida* recipient, which were maintained at 30 °C throughout.

For ACE experiments involving *E. coli Syn61* donors and for iterative ACE for trimera generation, donor and recipient cultures were inoculated into 10 mL of LB media with appropriate antibiotics for overnight growth (12-16 hrs). Overnight cultures were then pelleted by centrifugation and replenished with new 10 mL of LB media with the appropriate antibiotics and 0.5% (w/v) arabinose and incubated at 37°C for 2 hours. After induction, the remainder of this abbreviated ACE procedure is the same as described above. A comparison of the two described ACE methods is discussed in Supplemental Note 2.

Colonies were picked from selective post-ACE plates for picked and streaked out on fresh selective LB-agar plate for overnight incubation at 37°C. Colonies were picked from the streak-out plate and grown overnight in selective LB for long-term storage in 15% (v/v) glycerol at –80°C. A list of targets used in ACE experiments is given in Supplemental Table 4, a list of ACE experiments is given in Supplemental Table 5 and a list of ACE chimeric genomes (GEOs) made is given in Supplemental Table 6.

For GFP+ c.f.u. ACE efficiency plots and *in vivo* ACE specificity screens, colonies of the correct phenotypes were counted on post-ACE plates and tabulated in Supplemental Table 7.

### Phenotyping and Genotyping Analyses of Transposition

Single colonies of post-ACE were picked from selective LB-agar streak-out plates and resuspended in 40 uL of milli-Q H_2_O. For colony phenotyping assays, 3-4 uL of the resuspension is spotted onto LB-agar plates with the target antibiotic. For colony genotyping PCR, reactions were performed with 2X Red Taq Master Mix from Apex Bioresearch products with 1 uL of the resuspension used as template per 20 uL of reaction volume, which contains 0.2 uM of each primer. Cycling protocol starts with 2 minutes of initial cell lysis and denaturation at 98°C followed by 36 cycles of 95°C denaturation, 60-65°C primer annealing and 72°C extension (at 1 Kb/minute) with a final extension of 5 minutes at 72°C. For junction detection, primer pairs contained one donor-specific primer and one recipient-specific primer and were varied so that all integration orientation could be assayed for both the LE and RE transposon junctions. For internal region detection of the transferred sequence, primer pairs containing two donor-specific primers were used. PCR amplicons were resolved by 1% agarose gel electrophoresis using SYBR Safe staining for visualization.

All oligonucleotides used in analysis and validation of post-ACE bacteria are listed in Supplemental Table 3.

### Growth Experiments

Bacterial cultures were inoculated the night before in LB and appropriate antibiotics and grown overnight for 12-16 hours. Cultures were pelleted by centrifugation (4000 rpm for 10 minutes) and spent media was replaced with the same volume of fresh LB without antibiotics. Cultures were then diluted to 0.04 into 1 mL of fresh LB without antibiotics and 200 uL of the dilution was aliquoted onto a 96-well plate. Growth experiments were performed with three technical replicates per strain. Growth experiment was performed on a Tecan Spark^®^ Multimode Microplate Reader for 20 hours at 37°C with double orbital shaking at 108 rpm. OD_600_ readings were taken every 15 minutes. Data was truncated at 12 hours to account for evaporation of the wells causing disturbance of the OD_600_ readings late into the experiment.

### Confocal microscope imaging

Morphology and phenotype of chimeric strains were observed with a confocal laser scanning microscope (Zeiss LSM 800). Bacteria cells are seeded on the Luria Broth (LB) agarose gel pads and then mounted on the glass-bottom confocal dish. Phase contrast and fluorescence images were acquired using an oil immersion objective plan-apochromat 100x/1.4.

### Whole genome sequencing (WGS)

Bacterial cultures were grown in LB media supplemented with the appropriate antibiotics until confluent. Cultures were pelleted by centrifugation and genomic DNA (gDNA) was extracted from the pellets using DNeasy Blood & Tissue Kit (QIAGEN^™^ cat # 69504) per manufacturer instructions. Concentrations of gDNA were quantified with the Invitrogen Qubit^™^ 4 Fluorometer using the 1x dsDNA High Sensitivity (HS) assay kit (Thermo Fisher Scientific). For library preparation, Nextera^™^ DNA Flex Library Prep Kit (Illumina) was used per manufacturer instructions to tag, barcode, amplify and add index primers (i5 and i7) to the library. 200 ng of starting gDNA was used per barcoded library. Prepared libraries were quantified with the Qubit^™^ 4 Fluorometer before pooling 15 ng per barcoded library. Pooled library denaturation and flow cell loading was performed per manufacturer instructions. Sequencing was performed on an Illumina MiSeq using MiSeq reagent kit v3 (600-cycles) with 300 bp paired-end reads and automated demultiplexing and adapter trimming.

### Transposon sequencing (Tn-Seq)

Colonies on post-ACE selection plates were scraped into 4 mL of LB and grown in appropriate antibiotics for 4 hours before 1 mL of culture was pelleted by centrifugation. gDNA was extracted from the pelleted cell populations using DNeasy Blood & Tissue Kit (QIAGEN^™^ cat # 69504) per manufacturer instructions. Concentrations of gDNA were quantified with the Invitrogen Qubit^™^ 4 Fluorometer. For Tn-Seq library preparation, 200 ng of starting gDNA was tagmented and washed with Nextera^™^ DNA Flex Library Prep Kit as per manufacturer instructions. For library amplification and barcoding, custom index 2 (i5) oligonucleotides annealing to the RE of *Tn6677* were used instead of the standard i5 indices (see Supplemental Table 3). These custom i5 oligos were used alongside standard index 1 (i7) oligos provided Illumina with 15x cycles of library amplification PCR at 65°C annealing (all else according to the manufacturer). Barcoded libraries were cleaned up as per manufacturer instructions and Qubit^™^ 4 Fluorometer before pooling 30 ng per library. Pooled library denaturation and flow cell loading was performed per manufacturer instructions. Prior to sequencing, custom sequencing oligos was spiked into the standard sequencing oligo well in the reagent kit to enable targeted sequencing of enriched transposon sequences within the libraries. Sequencing was performed on an Illumina MiSeq using MiSeq reagent kit v3 (600-cycles) with 300 bp paired-end reads and automated demultiplexing and adapter trimming.

### RNA sequencing (RNA-Seq)

For RNA-Seq of *E. coli* DH10B Δ*dif* + 2-Mb Syn61 Syn-seg. 1 and its progenitor donor and recipient strains, bacterial cultures were grown in LB media supplemented with the appropriate antibiotics overnight. The next day, the cultures were back diluted to OD_600_ of 0.05 in fresh 10 mL of LB without any antibiotics and grown to an OD_600_ of 0.6. 1 mL of the culture was pelleted by centrifugation for RNA extraction using RNeasy Protect Bacteria Mini Kit (QIAGEN^™^ cat # 74524). Extracted RNA was quantified using Invitrogen Qubit^™^ 4 Fluorometer using the RNA High Sensitivity (HS) assay kit (Thermo Fisher Scientific). Sequencing libraries from extracted RNA samples were prepped and sequenced on an Illumina HiSeq with 50 bp paired-end reads by the Millard and Muriel Jacobs Genetics and Genomics Laboratory at the California Institute of Technology. RNA-Seq experiments were performed with three technical replicates per strain.

For RNA-Seq and differential gene expression of *E. coli* DH10B Δdif::Syn. Seg 1-*S. flexneri*-1Mb*dif, E. coli* DH10B *Δdif::S. flexneri*-1Mb*dif, E. coli* MDS42 *Δdif::S. flexneri*-1Mb*dif* and associated donor and recipient strains, and additionally for differential gene expression of *E. coli* DH10B Δ*dif* + 2-Mb Syn61 Syn-seg. 1 and associated donor and recipient strains, bacterial cultures were grown in LB media supplemented with the appropriate antibiotics overnight. The next day, the cultures were back diluted to OD_600_ of 0.05 in fresh 10 mL of LB without any antibiotics and grown to an OD_600_ of 0.6. 500 uL of the culture was pelleted by centrifugation and frozen for submission to GENEWIZ (USA; www.genewiz.com), for RNA extraction with rRNA depletion, library preparation and standard RNA sequencing on an Illumina NovaSeq with 150 bp paired-end reads. RNA-Seq experiments were performed with three technical replicates per strain.

### WGS analysis

Sequencing reads from whole-genome samples were aligned to expected reference genomes using the Bowtie2 short-read aligner^5^. Alignment files were further processed with Samtools^6^ and deepTools^7^ was used to estimate RPGC throughout the expected reference genome in 10-kb bins. References for gene homolog variants for the recipient and donor strains were identified via a reciprocal alignment approach using Blast^8^. Identified gene homolog variants were used as references to estimate, independent of short-read sequence alignment, the read fraction allocated to each respective reference position. The read fraction estimation per reference was calculated as the sum of counts across 14-bp bins encompassing distinct nucleotide polymorphisms detected via multiple-sequence alignment using MUSCLE^9^. A list of SRA accessions linked to all WGS data presented throughout this work can be found in Supplemental Table 8.

### Tn-Seq analysis

Custom Python pipelines were developed to process Tn-Seq data generated from ACE experiments, as described^10^ in prior work. Briefly, reads containing the 15-bp 5′-terminal sequence of the enriched Tn arm end were then selected and the upstream 17-bp region was used as a genomic footprint corresponding to the integration site in the genome. Fingerprint sequences were then aligned to expected reference genomes using Bowtie2. Alignment positions documented per read were later grouped into unique 10-kb bins, and unless stated otherwise insertion frequencies were plotted as a percentage of total mapping reads. The on-target percentage was calculated as the percentage of reads corresponding to integration events within the 10-kb window flanking the target integration site. The integration orientation bias is defined as the ratio of number of reads corresponding to T-RL insertions to those corresponding to T-LR insertions. A list of SRA accessions linked to all Tn-Seq data presented throughout this work can be found in Supplemental Table 9.

### RNA-Seq analysis

Custom Python pipelines were developed to quantify RNA sequencing reads uniquely mapping to gene homolog references. Sequencing reads from whole-transcriptome RNA samples were first aligned to expected transcriptome references per expected genome reference using the Bowtie2 short-read aligner. Because reads can map to multiple gene homolog variants, counts were allocated to specific gene homolog references by using discriminatory 14-bp bins per reference (identified via MUSCLE). The raw counts per gene homolog reference was then transformed to log(TPM+1) and plotted using various visualization and analysis pipelines written in Python. For differential expression analysis, raw counts of triplicate samples were further processed via DESeq2^11^, internally normalizing samples and calculating fold-change and adjusted p-values per transcript reference. DESeq2 results were illustrated as a volcano plot, highlighting enrichment of redundant genes within expected references. Gene ontology pathway enrichment analysis was performed using ShinyGO^12^ on the intersection of over-expressed genes in chimeric ACE samples, as defined by an adjusted p-value < 0.001 and a fold-change ≥ 1. A list of SRA accessions linked to all RNA-Seq data presented throughout this work can be found in Supplemental Table 10.

### Data reporting

No statistical methods were used to predetermine sample size. The experiments were not randomized, and investigators were not blinded to allocation during experiments and outcome assessment.

### Statistical analysis

Unless stated otherwise, statistical analysis on triplicates for experiments describing the efficiency and precision of ACE were performed by calculating the mean and standard deviation (SD) of each experimental sample group. For transcriptomic analysis, raw counts were converted to log(TPM+1), where each transcript count was normalized by the expected transcript reference length. Differential expression analysis was carried out using DESeq2^11^ using raw counts of samples as input. The following cutoffs were applied to demark significant changes in RNA expression in the context of massively expanded genomes: a base expression value (raw count) > 5, an adjusted p-value < 0.001, and a fold-change of 1. Gene ontology pathway enrichment analysis was performed using standard parameters in ShinyGO 0.77, only showing up to 10 pathways^12^.

---

**Supplemental Note 1**

Off-target patterns of mutant CAST variants during ACE transfers were interrogated with Tn-Seq. In the case of CAST-crRNA^NT^ transfer, a single position is enriched in mapped reads possibly due to partial complementarity of crRNA^NT^ with genomic sequences (**Extended Figure 2f**). Transfers with CAST-ΔcrRNA and CAST-Δcascade exhibited more genome-wide random transposition without inherent biases (**Extended Figure 2g-h**). However, the presence of a semi-enriched integration site on the genome rearises when TniQ is deleted on top of Δ*cascade* in Δ*Qcascade*. Target-selection studies of CASTs have uncovered multiple families employing alternative RNA-independent targeting pathways in addition to canonical and noncanonical crRNA-dependent transposition^1,2,3,4^. Whether Type I-F *TniQ-TnsABC* constitutes an RNA-independent transposition complex analogous to that of the Type V-K TniQ-TnsBC complex^5^, or whether the transposome *TnsABC* has intrinsic sequence specificity like Type I-B, Type I-F and I-D *TnsD* is subject to future investigation^2,4,6^.

**Supplemental Note 2**

The audience might observe differences between the ACE protocol used in figure 2 and figure 5 and that which is used in figure 3 and figure 4 (see methods). While both versions of the protocol feature a solid agar conjugation step followed by a recovery and outgrowth step before selective plating, figure 2 and figure 5 experiments were additionally performed with cultures grown to OD600 0.5 from a dilution of the overnight culture. However, this initial culturing step contributed a significant amount of additional time to complete the entire protocol, limiting experimental throughput and necessitating the work of two researchers to sustainably complete one experiment. Moreover, the donor and recipient strains used in the experiments typically had different doubling times, so it was challenging to coordinate the growth of both cultures to the OD600 0.5 growth endpoint. By replacing this initial growth step with induction of the overnight culture in fresh media, we can reduce the protocol time by an average of three hours while also decreasing handling volumes, enabling higher experimental throughput and less burden on individual researchers to reasonably carry out megabase-scale ACE in a single day. This reasoning is what led us to adopt the modified protocol for figure 3 experiments.

On top of these practical experimental considerations, we also observed an increase in experimental efficiency (measured by GFP+ c.f.u.) when carrying out the modified protocol compared to the new original protocol with 1-Mb Syn61 dif donor transfers with ACE (**Show figure**), possibly due to larger populations of donor and recipient bacteria in the overnight culture compared to daytime culture. We observe little to no instability when performing this protocol for generating *DH10B* + 1-Mb Syn61 and *DH10B* + 2-Mb Syn61 Syn-seg 1, leading us to adopt this protocol for experiments in figure 4 (figure 5 experiments were already done with the original figure 2 protocol). Shortening or outright eliminating the post conjugation recovery and outgrowth step and directly plating the conjugation spots on selective media further reduced of protocol time and was done for some of the extended figure experiments.

## References

1. Boeke, J. D. et al. The Genome Project-Write. Science 353, 126–127 (2016).

2. Ostrov, N. et al. Technological challenges and milestones for writing genomes. Science 366, 310–312 (2019).

3. Zhang, W., Mitchell, L. A., Bader, J. S. & Boeke, J. D. Synthetic Genomes. Annu. Rev. Biochem. 89, 77–101 (2020).

4. Jiang, S., Zhao, S., Cai, Z., Tang, Y. & Dai, J. Synthetic yeast genomes for studying chromosomal features. Current Opinion in Systems Biology 23, 1–7 (2020).

5. Coradini, A. L. V., Hull, C. B. & Ehrenreich, I. M. Building genomes to understand biology. Nat Commun 11, 6177 (2020).

6. Venter, J. C., Glass, J. I., Hutchison, C. A. & Vashee, S. Synthetic chromosomes, genomes, viruses, and cells. Cell 185, 2708–2724 (2022).

7. Hutchison, C. A. et al. Design and synthesis of a minimal bacterial genome. Science 351, aad6253 (2016).

8. Richardson, S. M. et al. Design of a synthetic yeast genome. Science 355, 1040–1044 (2017).

9. Wang, K. et al. Defining synonymous codon compression schemes by genome recoding. Nature 539, 59–64 (2016).

10. Shen, Y. et al. Deep functional analysis of synII, a 770-kilobase synthetic yeast chromosome. Science 355, eaaf4791 (2017).

11. Mercy, G. et al. 3D organization of synthetic and scrambled chromosomes. Science 355, eaaf4597 (2017).

12. Schindler, D. et al. Design, construction, and functional characterization of a tRNA neochromosome in yeast. Cell S0092867423011303 (2023) doi:10.1016/j.cell.2023.10.015.

13. Lajoie, M. J. et al. Genomically Recoded Organisms Expand Biological Functions. Science 342, 357–360 (2013).

14. Fredens, J. et al. Total synthesis of Escherichia coli with a recoded genome. Nature 569, 514–518 (2019).

15. Robertson, W. E. et al. Sense codon reassignment enables viral resistance and encoded polymer synthesis. Science 372, 1057–1062 (2021).

16. Nyerges, A. et al. A swapped genetic code prevents viral infections and gene transfer. Nature 615, 720–727 (2023).

17. Gibson, D. G. et al. Complete Chemical Synthesis, Assembly, and Cloning of a Mycoplasma genitalium Genome. Science 319, 1215–1220 (2008).

18. Gibson, D. G. et al. Creation of a Bacterial Cell Controlled by a Chemically Synthesized Genome. Science 329, 52–56 (2010).

19. Zhao, Y. et al. Debugging and consolidating multiple synthetic chromosomes reveals combinatorial genetic interactions. Cell S0092867423010796 (2023) doi:10.1016/j.cell.2023.09.025.

20. Ostrov, N. et al. Design, synthesis, and testing toward a 57-codon genome. Science 353, 819–822 (2016).

21. Lau, Y. H. et al. Large-scale recoding of a bacterial genome by iterative recombineering of synthetic DNA. Nucleic Acids Research 45, 6971–6980 (2017).

22. Venetz, J. E., et al. Chemical synthesis rewriting of a bacterial genome to achieve design flexibility and biological functionality. Proc. Natl. Acad. Sci. U.S.A. 116, 8070–8079 (2019).

23. Annaluru, N. et al. Total Synthesis of a Functional Designer Eukaryotic Chromosome. Science 344, 55–58 (2014).

24. Lartigue, C. et al. Creating Bacterial Strains from Genomes That Have Been Cloned and Engineered in Yeast. Science 325, 1693–1696 (2009).

25. Wang, K., De La Torre, D., Robertson, W. E. & Chin, J. W. Programmed chromosome fission and fusion enable precise large-scale genome rearrangement and assembly. Science 365, 922–926 (2019).

26. Yoneji, T., Fujita, H., Mukai, T. & Su’etsugu, M. Grand scale genome manipulation via chromosome swapping in *Escherichia coli* programmed by three one megabase chromosomes. Nucleic Acids Research 49, 8407–8418 (2021).

27. Isaacs, F. J. et al. Precise Manipulation of Chromosomes in Vivo Enables Genome-Wide Codon Replacement. Science 333, 348–353 (2011).

28. Zürcher, J. F. et al. Continuous synthesis of E. coli genome sections and Mb-scale human DNA assembly. Nature 619, 555–562 (2023).

29. Itaya, M., Tsuge, K., Koizumi, M. & Fujita, K. Combining two genomes in one cell: Stable cloning of the Synechocystis PCC6803 genome in the Bacillus subtilis 168 genome. Proc. Natl. Acad. Sci. U.S.A. 102, 15971–15976 (2005).

30. Mukai, T. et al. Overcoming the Challenges of Megabase-Sized Plasmid Construction in *Escherichia coli*. ACS Synth. Biol. 9, 1315–1327 (2020).

31. Strand, T. A., Lale, R., Degnes, K. F., Lando, M. & Valla, S. A New and Improved Host-Independent Plasmid System for RK2-Based Conjugal Transfer. PLoS ONE 9, e90372 (2014).

32. Klompe, S. E., Vo, P. L. H., Halpin-Healy, T. S. & Sternberg, S. H. Transposon-encoded CRISPR–Cas systems direct RNA-guided DNA integration. Nature 571, 219–225 (2019).

33. Vo, P. L. H. et al. CRISPR RNA-guided integrases for high-efficiency, multiplexed bacterial genome engineering. Nat Biotechnol 39, 480–489 (2021).

34. Rubin, B. E. et al. Species– and site-specific genome editing in complex bacterial communities. Nat Microbiol 7, 34–47 (2021).

35. Campbell, B. C., et al. mGreenLantern: a bright monomeric fluorescent protein with rapid expression and cell filling properties for neuronal imaging. Proc. Natl. Acad. Sci. U.S.A. 117, 30710–30721 (2020).

36. Virolle, C., Goldlust, K., Djermoun, S., Bigot, S. & Lesterlin, C. Plasmid Transfer by Conjugation in Gram-Negative Bacteria: From the Cellular to the Community Level. Genes 11, 1239 (2020).

37. Klompe, S. E. et al. Evolutionary and mechanistic diversity of Type I-F CRISPR-associated transposons. Molecular Cell 82, 616–628.e5 (2022).

38. Pósfai, G. et al. Emergent Properties of Reduced-Genome *Escherichia coli*. Science 312, 1044–1046 (2006).

39. Mulcair, M. D. et al. A Molecular Mousetrap Determines Polarity of Termination of DNA Replication in E. coli. Cell 125, 1309–1319 (2006).

40. Duggin, I. G. & Bell, S. D. Termination Structures in the Escherichia coli Chromosome Replication Fork Trap. Journal of Molecular Biology 387, 532–539 (2009).

41. Rudolph, C. J., Upton, A. L., Stockum, A., Nieduszynski, C. A. & Lloyd, R. G. Avoiding chromosome pathology when replication forks collide. Nature 500, 608–611 (2013).

42. Wendel, B. M., Courcelle, C. T. & Courcelle, J. Completion of DNA replication in Escherichia coli. Proc. Natl. Acad. Sci. U.S.A. 111, 16454–16459 (2014).

43. Bidnenko, V. Replication fork collapse at replication terminator sequences. The EMBO Journal 21, 3898–3907 (2002).

44. Esnault, E., Valens, M., Espéli, O. & Boccard, F. Chromosome Structuring Limits Genome Plasticity in Escherichia coli. PLoS Genet 3, e226 (2007).

45. Lesterlin, C., Pages, C., Dubarry, N., Dasgupta, S. & Cornet, F. Asymmetry of Chromosome Replichores Renders the DNA Translocase Activity of FtsK Essential for Cell Division and Cell Shape Maintenance in Escherichia coli. PLoS Genet 4, e1000288 (2008).

46. Marman, H. E., Mey, A. R. & Payne, S. M. Elongation Factor P and Modifying Enzyme PoxA Are Necessary for Virulence of Shigella flexneri. Infect Immun 82, 3612–3621 (2014).

47. Koestler, B. J. et al. Human Intestinal Enteroids as a Model System of *Shigella* Pathogenesis. Infect Immun 87, e00733–18 (2019).

48. Wei, J. et al. Complete Genome Sequence and Comparative Genomics of *Shigella flexneri* Serotype 2a Strain 2457T. Infect Immun 71, 2775–2786 (2003).

49. Weinstock, M. T., Hesek, E. D., Wilson, C. M. & Gibson, D. G. Vibrio natriegens as a fast-growing host for molecular biology. Nat Methods 13, 849–851 (2016).

50. Martínez-García, E., Nikel, P. I., Aparicio, T. & De Lorenzo, V. Pseudomonas 2.0: genetic upgrading of P. putida KT2440 as an enhanced host for heterologous gene expression. Microb Cell Fact 13, 159 (2014).

51. Dalia, T. N. et al. Multiplex Genome Editing by Natural Transformation (MuGENT) for Synthetic Biology in *Vibrio natriegens*. ACS Synth. Biol. 6, 1650–1655 (2017).

52. Thorpe, H. M. & Smith, M. C. M. In vitro site-specific integration of bacteriophage DNA catalyzed by a recombinase of the resolvase/invertase family. Proc. Natl. Acad. Sci. U.S.A. 95, 5505–5510 (1998).

53. Stellwagen, A. E. Avoiding self: two Tn7-encoded proteins mediate target immunity in Tn7 transposition. The EMBO Journal 16, 6823–6834 (1997).

54. Halpin-Healy, T. S., Klompe, S. E., Sternberg, S. H. & Fernández, I. S. Structural basis of DNA targeting by a transposon-encoded CRISPR–Cas system. Nature 577, 271–274 (2020).

55. Hoffmann, F. T. et al. Selective TnsC recruitment enhances the fidelity of RNA-guided transposition. Nature 609, 384–393 (2022).

56. Park, J.-U. et al. Multiple adaptations underly co-option of a CRISPR surveillance complex for RNA-guided DNA transposition. Molecular Cell 83, 1827–1838.e6 (2023).

57. Wimmer, F., Mougiakos, I., Englert, F. & Beisel, C. L. Rapid cell-free characterization of multi-subunit CRISPR effectors and transposons. Molecular Cell 82, 1210–1224.e6 (2022).

58. Lampe, G. D. et al. Targeted DNA integration in human cells without double-strand breaks using CRISPR-associated transposases. Nat Biotechnol (2023) doi:10.1038/s41587-023-01748-1.

59. Strecker, J. et al. RNA-guided DNA insertion with CRISPR-associated transposases. Science 365, 48–53 (2019).

60. Vo, P. L. H., Acree, C., Smith, M. L. & Sternberg, S. H. Unbiased profiling of CRISPR RNA-guided transposition products by long-read sequencing. Mobile DNA 12, 13 (2021).

61. Tou, C. J., Orr, B. & Kleinstiver, B. P. Precise cut-and-paste DNA insertion using engineered type V-K CRISPR-associated transposases. Nat Biotechnol 41, 968–979 (2023).

62. George, J. T. et al. Mechanism of target site selection by type V-K CRISPR-associated transposases. Science 382, eadj8543 (2023).

63. Fu, J. et al. Efficient transfer of two large secondary metabolite pathway gene clusters into heterologous hosts by transposition. Nucleic Acids Research 36, e113–e113 (2008).

64. Huo, L. et al. Heterologous expression of bacterial natural product biosynthetic pathways. Nat. Prod. Rep. 36, 1412–1436 (2019).

65. Ryu, M.-H. et al. Control of nitrogen fixation in bacteria that associate with cereals. Nat Microbiol 5, 314–330 (2019).

66. Scherlach, K. & Hertweck, C. Mining and unearthing hidden biosynthetic potential. Nat Commun 12, 3864 (2021).

## References

1. Wang, M., Sanfiorenzo, C., Zhang, R. J. & Wang, K. A universal system for streamlined genome integrations with CRISPR-associated transposases. bioRxiv 2022.05.30.494051 (2022) doi:10.1101/2022.05.30.494051.

2. Weinstock, M. T., Hesek, E. D., Wilson, C. M. & Gibson, D. G. Vibrio natriegens as a fast-growing host for molecular biology. Nat Methods 13, 849–851 (2016).

3. Yu, D., et al. An efficient recombination system for chromosome engineering in Escherichia coli. Proc. Natl. Acad. Sci. U.S.A. 97, 5978–5983 (2000).

4. Datsenko, K. A. & Wanner, B. L. One-step inactivation of chromosomal genes in Escherichia coli K-12 using PCR products. Proc. Natl. Acad. Sci. U.S.A. 97, 6640–6645 (2000).

5. Langmead, B. & Salzberg, S. L. Fast gapped-read alignment with Bowtie 2. Nat Methods 9, 357–359 (2012).

6. Li, H. et al. The Sequence Alignment/Map format and SAMtools. Bioinformatics 25, 2078– 2079 (2009).

7. Ramírez, F., Dündar, F., Diehl, S., Grüning, B. A. & Manke, T. deepTools: a flexible platform for exploring deep-sequencing data. Nucleic Acids Research 42, W187–W191 (2014).

8. Altschul, S. F., Gish, W., Miller, W., Myers, E. W. & Lipman, D. J. Basic local alignment search tool. Journal of Molecular Biology 215, 403–410 (1990).

9. Edgar, R. C. MUSCLE: a multiple sequence alignment method with reduced time and space complexity. BMC Bioinformatics 5, 113 (2004).

10. Vo, P. L. H. et al. CRISPR RNA-guided integrases for high-efficiency, multiplexed bacterial genome engineering. Nat Biotechnol 39, 480–489 (2021).

11. Love, M. I., Huber, W. & Anders, S. Moderated estimation of fold change and dispersion for RNA-seq data with DESeq2. Genome Biol 15, 550 (2014).

12. Ge, S. X., Jung, D. & Yao, R. ShinyGO: a graphical gene-set enrichment tool for animals and plants. Bioinformatics 36, 2628–2629 (2020).

## References

1. Petassi, M. T., Hsieh, S.-C. & Peters, J. E. Guide RNA Categorization Enables Target Site Choice in Tn7-CRISPR-Cas Transposons. Cell 183, 1757–1771.e18 (2020).

2. Saito, M. et al. Dual modes of CRISPR-associated transposon homing. Cell 184, 2441–2453.e18 (2021).

3. Faure, G. et al. Modularity and diversity of target selectors in Tn7 transposons. Mol. Cell 83, 2122–2136.e10 (2023).

4. Klompe, S. E. et al. Evolutionary and mechanistic diversity of Type I-F CRISPR-associated transposons. Mol. Cell 82, 616–628.e5 (2022).

5. George, J. T. et al. Mechanism of target site selection by type V-K CRISPR-associated transposases. Science 382, eadj8543 (2023).

6. Hsieh, S.-C. & Peters, J. E. Discovery and characterization of novel type I-D CRISPR-guided transposons identified among diverse Tn7-like elements in cyanobacteria. Nucleic Acids Res. 51, 765–782 (2023).

